# Stepwise expansion of recombination suppression on sex chromosomes and other supergenes through lower load advantage and deleterious mutation sheltering

**DOI:** 10.1101/2025.06.27.661902

**Authors:** Paul Jay, Amandine Véber, Tatiana Giraud

## Abstract

Many organisms possess sex chromosomes with non-recombining regions that have expanded progressively. Yet, the causes of this stepwise expansion remain poorly understood. Here, using mathematical modeling and stochastic simulations, we show that the widespread presence of deleterious recessive mutations in genomes can generate several phenomena that can contribute to the evolution of recombination suppression on sex chromosomes. We demonstrate that a significant proportion of new chromosomal inversions, or any other recombination suppressor, are initially advantageous because they carry fewer mutations than average. However, these less-loaded inversions generally fail to fix on autosomes because, as their frequency increases, the recessive deleterious mutations they carry are more likely to occur in a homozygous state, leading to a selective disadvantage. In contrast, the permanent heterozygosity of Y-like sex chromosomes shelters their inversions from this disadvantage, facilitating their fixation and thereby the stepwise expansion of non-recombining regions. Due to this sheltering effect, we show that fixation probabilities are substantially higher on Y-like chromosomes than on autosomes across a wide range of parameters, particularly when segregating deleterious mutations are moderately to strongly recessive, and this is true even after controlling for differences in effective population sizes between those chromosomes. Then, once recombination is suppressed, deleterious mutations accumulate on inversions, which could select for recombination restoration. However, we show that the accumulation of overlapping genomic rearrangements following recombination suppression can prevent its restoration. Overall, our theoretical model proposes a testable framework that may contribute to explaining evolutionary strata on sex and mating-type chromosomes, and other supergenes.

## Introduction

Many organisms have sex chromosomes with large non-recombining regions that expand in a stepwise manner, although the underlying mechanisms remain poorly understood (Jay et al., 2024; Ponnikas et al., 2018; Saunders & Muyle, 2024; Wright et al., 2016). It has long been considered that recombination suppression on sex chromosomes gradually expands because selection favors the linkage of sexually antagonistic loci to sex-determining genes (Rice, 1987; Ruzicka et al., 2020; Wright et al., 2016), generating evolutionary strata of differentiation between sex chromosomes. However, there has been no compelling evidence that sexual antagonism is actually responsible for the expansion of recombination suppression in sex chromosome so far (Beukeboom & Perrin, 2014; Ironside, 2010; Jay et al., 2024; Ponnikas et al., 2018 but see Wright et al., 2017) and recombination suppression has also been reported to gradually expand around many fungal mating-type loci and other supergenes despite the lack of sexual antagonism (Bazzicalupo et al., 2019; Branco et al., 2017; Hartmann et al., 2021; Jay et al., 2021, 2024; Wang et al., 2013; Yan et al., 2020). In addition, theoretical issues have been raised about the model of sexual antagonism driving the evolution of sex chromosomes (Cavoto et al., 2018). Altogether, this suggests that other mechanisms can drive the stepwise extension of recombination suppression (Jay et al., 2024).

In 2021, we developed a model to explain the evolution of recombination suppression on sex chromosomes and other supergenes without assuming any form of sexual antagonism. It was based on the observation that most mutations occurring in genome are deleterious and partially recessive, resulting in a genomic landscape with many deleterious recessive mutations segregating at low frequencies within populations (Agrawal & Whitlock, 2011, 2012; Eyre-Walker & Keightley, 2007). The present paper is a revised version of our previous study (Jay et al., 2022), presenting new analyses of our original derivations and simulations, along with new simulations incorporating additional controls, which further support our theory and address previous criticisms (Charlesworth & Olito, 2024; Lenormand & Roze, 2024; Olito & Charlesworth, 2023). The rationale behind our theory is illustrated in Figure 1 and consists of three steps that could explain the evolution and maintenance of recombination suppression on sex chromosomes under some conditions. These steps are independent and can each contribute separately or in combination to the evolution of sex chromosomes.

**Figure 1.**
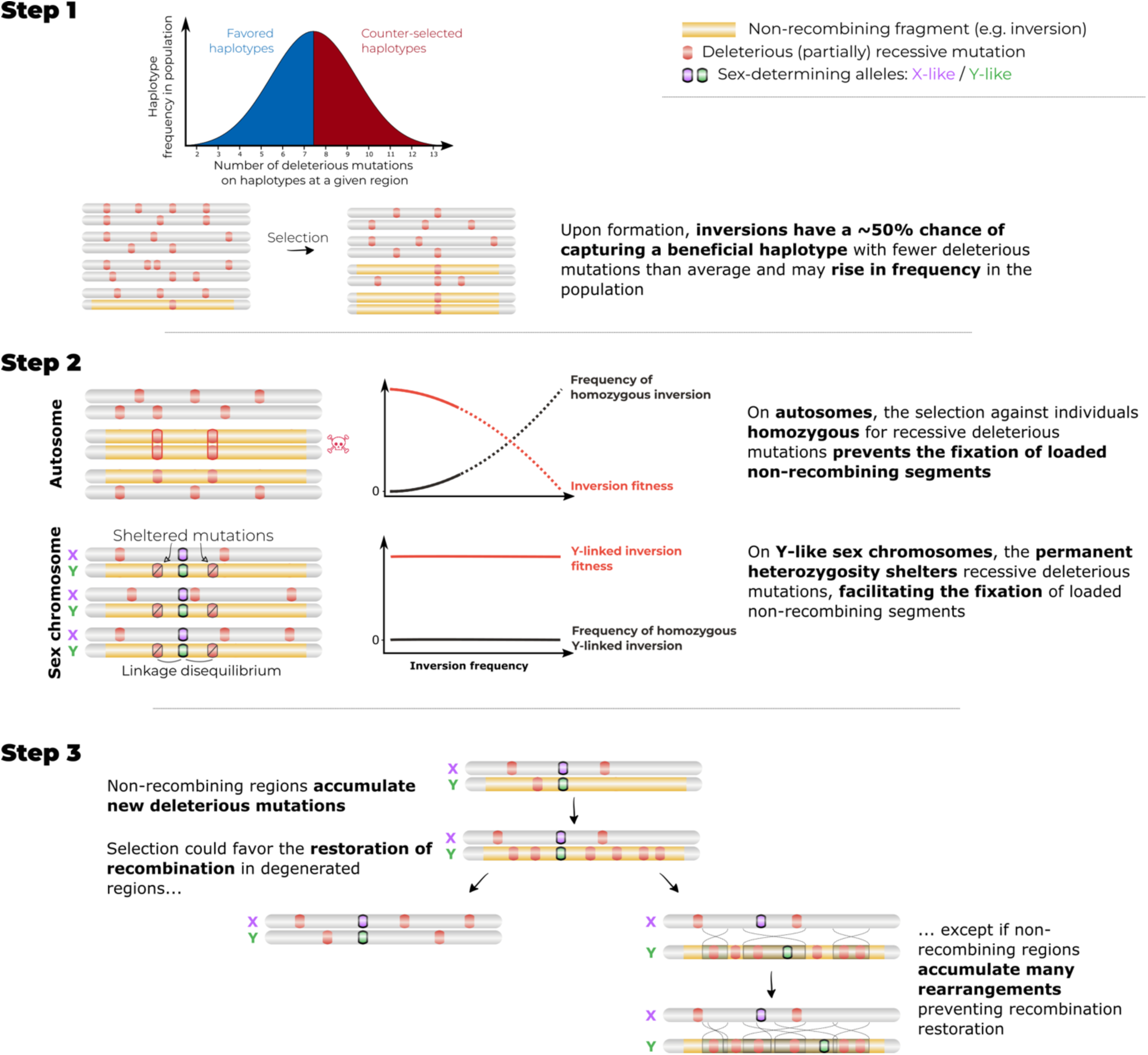
Schematic diagram of the three steps of the model. **Step 1.** Within any population and for any genomic region, diploid individuals (here represented by two homologous chromosomes) and haplotypes (i.e., specific combinations of mutations) carry variable numbers of partially recessive, deleterious mutations. A substantial fraction (about 50%) of haplotypes has fewer deleterious mutations than average in the corresponding DNA fragment and should be favored by selection. Chromosomal inversions therefore have a significant chance of capturing beneficial haplotypes. This is the lower-load advantage. **Step 2.** On autosomes, the increase in frequency of a beneficial inversion leads to a higher frequency of homozygotes for the inversion, which have a fitness disadvantage because they are homozygous for all the recessive deleterious mutations carried by the inversion. The mean fitness of the inversion in the population therefore decreases with increasing inversion frequency, preventing the inversion to reach a high frequency. Permanent heterozygosity at a Y-like sex-determining allele protects linked inversions from this homozygosity disadvantage. Hence, beneficial inversions (e.g., carrying fewer deleterious mutations than average) can continue increasing in frequency and become fixed in the population of Y-like chromosomes. This is the sheltering effect. **Step 3.** Fixed inversions on Y-like chromosomes suppress recombination within the corresponding segment. As a result of selective interference and Muller’s ratchet-like processes, the non-recombining region accumulates new deleterious mutations. This degeneration should favor the restoration of recombination, for example through inversion reversion. However, the accumulation of additional chromosomal rearrangements during degeneration may prevent the restoration of recombination.

The first step in our model corresponds to the selection of inversions (or any recombination suppressors) carrying by chance a lower load than average in the genomic region (Figure 1). As previously highlighted (Nei et al., 1967), the stochastic distribution of deleterious mutations generates variation in fitness at any genomic region due to differences in mutation load. Chromosomal inversions occurring at random in genomes will capture a specific haplotype of deleterious variants that can have a higher or lower fitness than average, just by chance. Inversions that capture fewer deleterious mutations than the population average for a given region (*i.e.*, have a lower mutation load) benefit from a relative fitness advantage and should therefore increase in frequency (Figure 1). While the inversions themselves are neutral in this hypothesis, their association with a favorable haplotype provides a selective benefit. As inversions suppress recombination when heterozygous, inverted advantageous haplotypes cannot recombine with less advantageous haplotypes and therefore keep their advantage through meiosis, in contrast to recombining haplotypes. Note that we will use the term “inversions” for simplicity, but the processes described in this paper hold with other mechanisms suppressing recombination in *cis*, such as epigenetic marks, regardless of whether they act only in heterozygotes, such as inversions, or also in homozygotes.

The second step in our model corresponds to the sheltering of the recessive mutations in the inversion due to their linkage with a permanently heterozygous allele, allowing inversions to fix despite their recessive load. Indeed, when a given inversion increases in frequency, homozygotes for the inversion also become more frequent. Provided that the inversion carries at least one recessive deleterious mutation, homozygotes will experience a disadvantage due to the expression of this recessive load. Such selection against homozygotes should prevent this inversion from reaching high frequencies. In contrast, if a loaded inversion captures a permanently heterozygous allele, such as a Y-like sex-determining gene, its deleterious mutations are maintained in a heterozygous state, preventing the expression of its recessive load and thereby facilitating its fixation on the Y-like chromosome. This protection against recessive load thanks to the linkage to a permanently heterozygous allele is hereafter called the sheltering effect. The successive fixation of additional inversions linked to this Y-fixed inversion by the same process should cause the non-recombining region to expand further, thereby leading to the formation of sex chromosomes with evolutionary strata.

It is important to note that the lower-load advantage (step 1) and the sheltering effect (step 2) are two distinct selective effects. The lower load is an intrinsic fitness advantage, driving the increase in frequency of inversions with fewer deleterious mutations than average; once the frequency of the less-loaded inversion has reached an appreciable level, a second phase starts during which the sheltering effect protects inversions linked to permanently heterozygous alleles from exposing their load at a homozygous state, allowing them to reach fixation. The sheltering effect could act in combination with other types of intrinsic advantage of inversions, such as sexual antagonism or local adaptation, provided that the inversions harbour at least one deleterious, partially recessive mutation (Jay et al., 2024).

The third step of our model corresponds to the maintenance of recombination suppression despite further mutation accumulation in less-loaded inversions. Once recombination is suppressed, the inversions will indeed accumulate deleterious mutations by selective interference and Muller’s ratchet-like processes (Bachtrog, 2013; Berdan et al., 2021a, 2023). The resulting degeneration may eventually select for recombination restoration, for instance *via* inversion reversion (Lenormand & Roze, 2022). However, the accumulation of overlapping rearrangements in such a region, due to genetic drift or selective mechanisms (Berdan et al., 2023), may hinder recombination restoration. Indeed, in such cases, reversion of the initial inversions would not restore collinearity and therefore recombination.

Here, we analyse the different mechanisms acting in the three steps described above, and show that, in combination, they can contribute to the evolution and maintenance of recombination suppression on sex chromosomes. The accumulation of deleterious mutations following recombination suppression has been extensively studied (Bachtrog, 2013; Blaser et al., 2014; Grossen et al., 2012; Nei, 1970), but we investigate here the converse, *i.e.*, that deleterious mutations could be a factor contributing to the evolution of recombination suppression, and not only a consequence.

An important point that has been subject to debate in the previous version of this paper is the right control to evaluate the efficacy of the sheltering effect (Charlesworth & Olito, 2024; Olito & Charlesworth, 2023; https://hal.science/hal-04763742). To assess whether the sheltering effect contributed to inversion fixation on the Y chromosome, we previously compared inversion fixation probabilities between autosomes (where the sheltering does not act) and the Y chromosome (where the sheltering acts), both with deleterious mutations segregating (Jay et al., 2022). It has been argued that inversion fixation probabilities should instead be compared between Y chromosomes in situations with versus without deleterious mutations segregating (Charlesworth & Olito, 2024; Olito & Charlesworth, 2023). However, this so-called “neutral” control (*i.e.*, Y chromosomes without any deleterious mutations) addresses a question different from that of the contribution of the sheltering effect in the fixation of inversions on the Y chromosome in natural populations: it tests whether the presence of deleterious mutations in genomes increases inversion fixation probabilities compared to a situation without any deleterious mutations in genomes. To assess the efficacy of the sheltering effect (*i.e.,* the protection from recessive deleterious mutations), one must compare situations where deleterious mutations are present, but with different levels of protection: autosomes (no sheltering) *versus* the Y chromosome (with sheltering). This is analogous to testing a drug: its efficacy is evaluated by comparing conditions with and without the protective drug, with the deleterious pathogen present in both cases. One would not assess the drug efficacy by comparing situations with and without the deleterious pathogen. The logic behind the different types of control and what they can show or cannot show is explained in more details in Box 1.

Here, to assess the contribution of the sheltering effect to the fixation of inversions on sex-chromosomes, we slightly changed our approach compared to our previous study (Jay et al., 2022) in order to take the difference in genetic drift between autosomes and Y chromosome into account (see Box 1): we now compare normalized fixation probabilities on autosomes and on Y chromosomes, dividing absolute fixation probabilities by the neutral fixation probabilities for each type of chromosome. Because neutral fixation probabilities depend directly and only on the effective population size, this normalization controls for differences in effective sizes between autosomes and the Y chromosome. Note that a recent study uses a similar approach (Roze & Lenormand, 2025). This normalization allows us to address simultaneously, but separately, two questions that were previously conflated: (i) the efficacy of the protective sheltering effect, and (ii) whether the presence of deleterious mutations in genomes increases inversion fixation probabilities. Following a previous criticism, we also now analyze inversion fixation probabilities from generation 1 (and not from generation 20 as in Jay et al. 2022) and show that this does not change the conclusions of Jay et al (2022).

##### What control should we use to assess the contribution of the lower-load advantage and of the sheltering effect to inversion fixation probabilities?

###### a) Evaluating the sheltering effect efficacy

To test the protective effect of a permanently heterozygous locus against the homozygous disadvantage incurred by inversions loaded with recessive deleterious mutations, one must evaluate how well inversions perform when they are linked to such a locus compared with when they are not, all else being equal (*i.e.*, identical population sizes and identical deleterious mutation parameters). As an analogy, consider evaluating the effect of a drug on the probability of developing a disease caused by pathogenic bacteria. Comparing the survival probability of individuals taking the drug in the presence of bacteria to those also taking the drug but without bacteria provides no information about the efficacy of the drug against the bacterial disease. To assess the benefit of the drug, one must compare survival between individuals taking the drug and those not taking it, in the presence of bacteria. In our case, recessive deleterious mutations play the role of the bacteria, potentially generating a fitness cost through homozygosity (the disease), which may be alleviated when the inversion becomes associated with a permanently heterozygous locus (this association playing the role of the drug). To evaluate the efficacy of the sheltering effect, we must therefore compare inversions with the protecting sheltering effect (on the Y chromosome) to those without the sheltering effect (on autosomes), in the presence of deleterious mutations.

However, an inversion cannot fix (in the classical population genetics sense, reaching 100% in the population) at a locus that is permanently heterozygous, but can only reach the frequency at which it becomes fully associated with one of the alleles (for example 25% if the locus corresponds to the male-determining allele on a Y chromosome). Consequently, direct comparisons between the probability of fixation of inversions in an autosome and the probability of becoming fully associated with a permanently heterozygous allele in a population of the same size (as in Jay et al 2022) are not fully appropriate to quantify the contribution of the sheltering effect to inversion fixation. Indeed, in the former case, the inversion must spread through a larger fraction of the population than in the latter. The effective population size experienced by the inversion, and thus genetic drift, therefore differs between the two situations.

Two possible approaches can be used to address this issue. First, one can ensure that the same effective population size is considered in both cases by adjusting census population sizes to compensate for the constraint imposed by permanent heterozygosity (for example, by simulating inversions on the Y chromosome in populations four times larger than those used for autosomal simulations). Second, one can normalize inversion fixation probabilities by the neutral fixation probability (the inverse of the effective population size) before comparing scenarios with and without permanently heterozygous loci. Because the probability of neutral fixation is dictated exclusively by effective population size, this normalization compensates for the disparity in effective sizes between autosomes and the Y chromosome. In this manuscript, we mainly adopt the second approach (but both approaches are used in Figure S7).

###### (b) Assessing the less-loaded advantage

Evaluating whether carrying a lower mutational load than average provides a selective advantage to inversions requires comparing the fixation probability of inversions with a lower load to that of inversions without such an advantage, all else being equal. Consequently, fixation probabilities cannot be directly compared between scenarios with and without deleterious mutations, because the conditions differ: the presence of mutations distorts the fitness landscape, it generates fitness variance within the population and temporal fluctuations in fitness that are absent in mutation-free simulations.

Several types of comparisons can instead be used to assess the lower-load advantage. One approach is to compare the fixation probability of inversions carrying fewer deleterious mutations than the population average with that of inversions carrying an average load. Alternatively, as done in the present study (Figure 2c), one can test whether the inversions that eventually reached fixation were those initially carrying the fewest deleterious mutations among all the randomly generated inversions.

**Figure 2.**
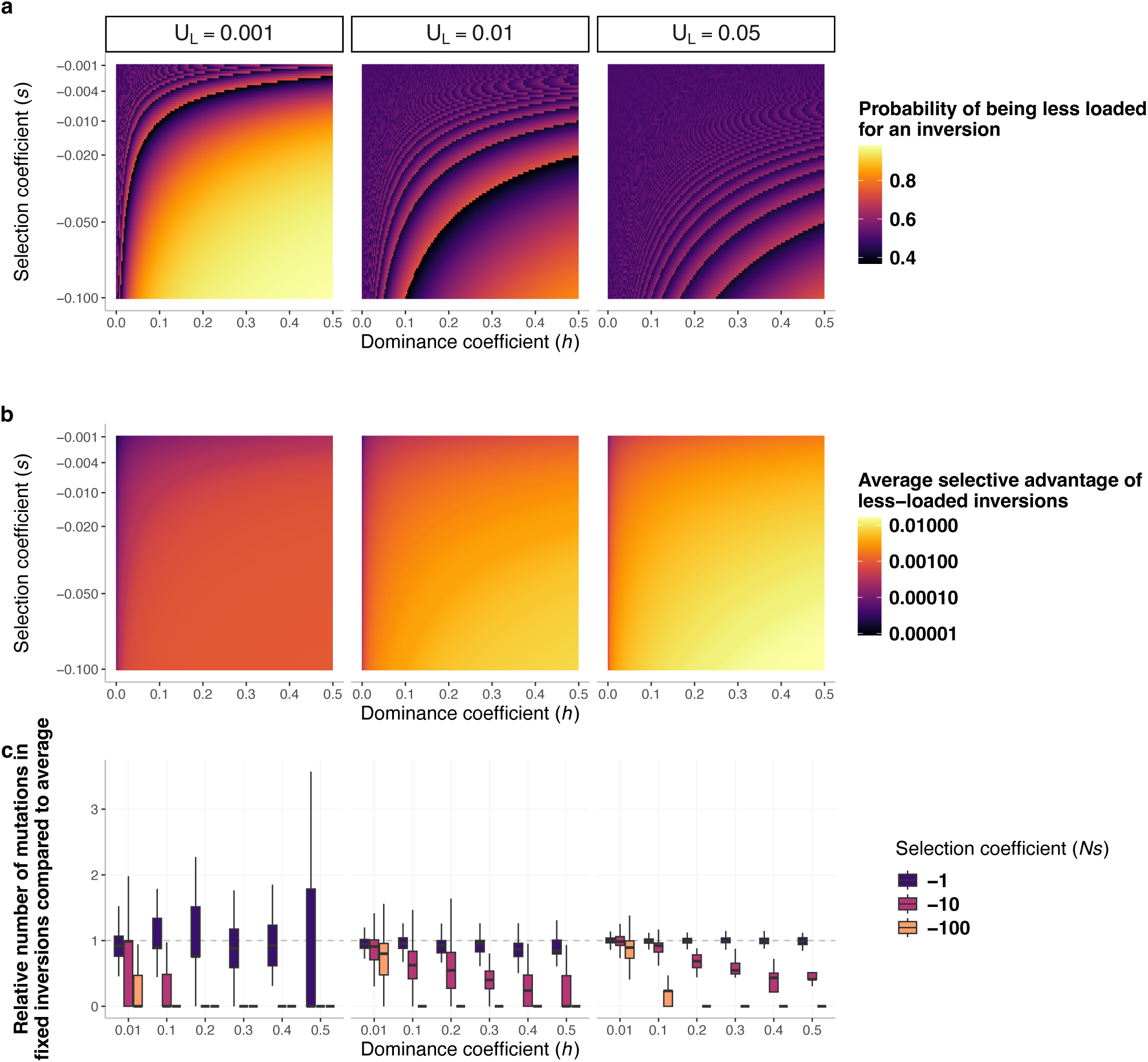
Less-loaded inversions are common and confer a fitness advantage. **a.** Probability that occurring inversions carry fewer mutations than average in an infinite population, as a function of the region-wide mutation rate (U_L_), the selection coefficient (s), and the dominance coefficient (h) of segregating mutations. Mutations are assumed to be at mutation–selection equilibrium. An inversion is considered “less-loaded” if the number m of mutations that it has captured is smaller than the average number of mutations, nq, obtained as the product of the size n of the inversion and of the equilibrium frequency of deleterious alleles q. As m is an integer, transitions between different values of m result in discontinuities in the plotted probabilities. **b.** Average initial selective advantage of less-loaded inversions in an infinite population. For each parameter combination, we averaged the fitness advantage of all inversions satisfying m < nq. The selective advantage of a given inversion was calculated by comparing the fitness of heterozygous carriers to the population mean fitness. On autosomes, the selective advantage of inversions should erode when the inversions increase in frequency because of the increasing frequency of individuals homozygous for the inversions, exposing the recessive load of these inversions. **c.** Relative number of mutations initially present in the inversions that eventually fixed on sex chromosomes in finite populations (N = 1000), normalized by the average number of mutations in non-inverted segments. Values below 1 indicate that fixed inversions initially had a lower load than average. The scripts used to produce this figure are available on GitHub. UL=0.001 corresponds to situation with u=1e-9 and n=1,000,000, whereas U_L_ values of 0.01 and 0.02 correspond to situations with u=1e-8 and n=1,000,000 or n=5,000,000, respectively.

###### (c) Comparing inversion probabilities on the Y chromosome in the presence and in the absence of deleterious mutations

The comparison between inversion fixation probabilities on Y-like chromosomes in the presence and in the absence of deleterious recessive mutations (the so-called “neutral case”) cannot be used to assess the contributions of the sheltering effect and the lower-load advantage to inversion fixation. Such a comparison only indicates whether the presence of deleterious mutations in the genome tends to favor or disfavor inversion fixation compared to the hypothetical scenario where all mutations are neutral. It does not allow one to perform model selection, for instance by arguing that a higher inversion fixation probability in the neutral case than in the case with deleterious recessive mutations implies that the neutral scenario better explains inversion fixation events observed in natural populations. Indeed, even if the fixation probability in scenario B is higher than in scenario A, determining the most likely cause of a fixation event observed in nature also requires considering the relative frequency with which each scenario occurs. For example, if the fixation probability in the neutral case (scenario B) is ten times higher than in the deleterious mutation case (scenario A), but genomes are 10^3 times more likely to contain deleterious recessive mutations than to be mutation-free, then the deleterious mutation scenario remains 100 times more likely to explain the observed inversion fixations.

## Method overview

We explored our three-step model of stepwise recombination suppression around permanently heterozygous alleles (Figure 1) with infinite population deterministic models and individual-based simulations. In both cases, we modeled diploid populations with only partially recessive deleterious mutations. Each mutation altered fitness by a factor of 1 + *hs* in heterozygotes and 1 + *s* in homozygotes, with *s*<0 and effects combining multiplicatively across loci. We first considered that all mutations occurring in genomes had the same dominance (*h*) and selection (*s*) coefficients and then relaxed this hypothesis. As the effect studied depends on the number of deleterious mutations in the studied region, we present our results as a function of *U_L_*, the region-wide deleterious mutation rate, defined as *U_L_* = *n*·*μ*, where *n* is the size of the region in base pairs and *μ* the per-site mutation rate. For reference, the estimated genome-wide deleterious mutation rates in humans and *Drosophila melanogaster* are approximately 2.2 and 1.3 mutations per genome per generation, respectively (Agrawal & Whitlock, 2012). Given the genome sizes of these species, an *n*=10 Mb region corresponds to *U_L_* ≈ 0.007 in humans and *U_L_* ≈ 0.07 in *D. melanogaster.* For simulations, we present the results as a function of the product *Ns*, where *N* is the census population size, because the evolutionary effect of selection depends on its strength relative to genetic drift.

To incorporate mutation dynamics during the spread of inversions within populations, previous studies have relied on mathematical models in which inversion dynamics are treated stochastically while mutation dynamics within the inversion are modeled deterministically (Olito et al. 2025; Olito & Abbott, 2024). This approach reflects the difficulty of incorporating into analytical models the stochastic behavior of numerous mutations and their selective interference, both within and between recombining and non-recombining haplotypes, particularly when the inversion frequency is changing through time. However, we argue that genetic drift and selective interference induced by genetic linkage are central to the evolution of inversions. For example, even in large populations, a newly arisen non-recombining segment initially has a very small effective population size. This implies that its mutational load –and thus its fitness trajectory– is strongly influenced by drift, at least during the early stages of its spread. Similarly, when an inversion increases in frequency, the effective population size of the non-inverted haplotypes is reduced, which in turn affects the accumulation of deleterious mutations in both inverted and non-inverted genomic segments. Generally, neglecting selective interference in the study of recombination suppression, while this is one of the processes thought to promote the evolution of recombination (Barton & Otto, 2005; Roze 2021), does not seem appropriate. Therefore, in this study we mostly used individual-based simulations to dissect the dynamics of inversions on autosomes and on Y-like sex chromosomes under realistic scenarios.

Individuals were considered to have two pairs of chromosomes, one of which harbored a locus with at least one allele permanently or almost permanently heterozygous (see Methods for details). Several situations were considered, mimicking those encountered in XY sex-determination systems, in fungal mating-type systems or in overdominant supergenes. We simulated the evolution of recombination modifiers suppressing recombination across the fragment in which they reside (*i.e*., *cis-*modifiers), either exclusively in heterozygotes (mimicking for example chromosomal inversions), or in both heterozygotes and homozygotes (*e.g*., histone modifications). Each of these recombination modifiers appeared in a single haplotype (*i.e.,* in a single chromosome of a single individual), and was thus in linkage disequilibrium with a specific set of mutations. We first considered recombination modifiers that were neutral by themselves, so that their fitness was exclusively dependent on the number of deleterious alleles within the captured segment. Therefore, the fate of these inversions was only influenced by their load (step 1) and possibly a sheltering effect (step 2). We then considered other scenarios, under which recombination modifiers were intrinsically beneficial. We first compared the dynamics of inversion-mimicking mutations in an autosome to those capturing a male-determining allele in an XY system, males being XY and females XX, the male-determining allele being permanently heterozygous; we then considered other types of recombination modifiers and heterozygosity rules. Due to computing limitations, only relatively small population sizes, *i.e.* of *N*=1,000 or *N*=10,000, were simulated.

## Results

### Less-loaded inversions are frequent in genomes and are advantageous (STEP 1)

As noted by several authors (Connallon & Olito, 2021; Nei et al., 1967; Olito & Abbott, 2020), inversions capturing fewer deleterious mutations than the population average in the focal genomic segment should be common in genomes and should increase in frequency. This can easily be verified by computing inversion fitness advantage as function of the point mutation rate (*u*), the mutation dominance coefficient (*h*), the mutation selective coefficient (*s*), the size of the inversion (*n*) and the number of mutations it carries (*m*) [see Supplementary text S1; (28, 37)]. Basic arithmetic shows that, under realistic parameter values, the vast majority of large chromosomal regions carry several deleterious mutations. For instance, considering *s*=-0.001, *h*=0.1, *μ*=1×10^-9^, more than 99.999% of 1Mb chromosomal fragments carry at least one mutation, the mean number of mutations being 10. In this context, inversions with fewer mutations than the population average –hereafter referred to as “less-loaded inversions”– tend to be common (Figure 2a). Indeed, when the region mutation rate *U_L_* is high, the distribution of mutation numbers across individuals is almost symmetric, so inversions have a 50% chance of being less-loaded. In contrast, when *U_L_* is low, this distribution is zero-inflated, increasing the probability of inversions being less-loaded over 50%. For instance, considering *U_L_*=0.001 (*e.g*., with *n*=1Mb and *μ*=1e-9), with *h* values ranging uniformly from 0 to 0.5 and *s* values ranging uniformly from −0.001 to −0.1, ca. 83% of the inversions carry fewer recessive deleterious mutations than average (Figure 2a).

The selective advantage of less-loaded inversions depends on the number of deleterious mutations they carry relative to the population average, as well as on the selection coefficients and dominance coefficients of those mutations. (Figure 2b). This advantage can be substantial, being for instance 0.0008 on average when *U*=0.001 and 0.0066 when *U*=0.05 under the range of parameter values studied (Figure 2b). Interestingly, the average selective advantage of less-loaded inversion decreases only slightly with decreasing mutation deleteriousness because such conditions lead to larger number of segregating mutations. Simulations of finite populations confirmed that the lower-load advantage could be a substantial driver of the increase in frequency of inversions, notably showing that inversions that went to fixation tended to initially harbour fewer mutations than average (Figure 2c). Large inversions (*i.e*., with higher *U_L_*), or inversions arising in mutation-dense regions, exhibit a greater fitness variance than small inversions, and therefore a potential greater selective advantage. As a consequence, the probability of fixation of Y-linked and autosomal inversions increased with both the inversion size and the mutation rate (Figure 3, 4 and S4, 5, 7). This indicates that the faster accumulation of deleterious mutations in larger inversions –due to their larger number of mutable sites– did not offset their larger initial selective advantage conferred by the lower-load effect. This contrasts with previous conclusions (Olito & Abbott, 2024), most likely because of differences in how mutation accumulation was modeled, or in the population sizes considered.

**Figure 3.**
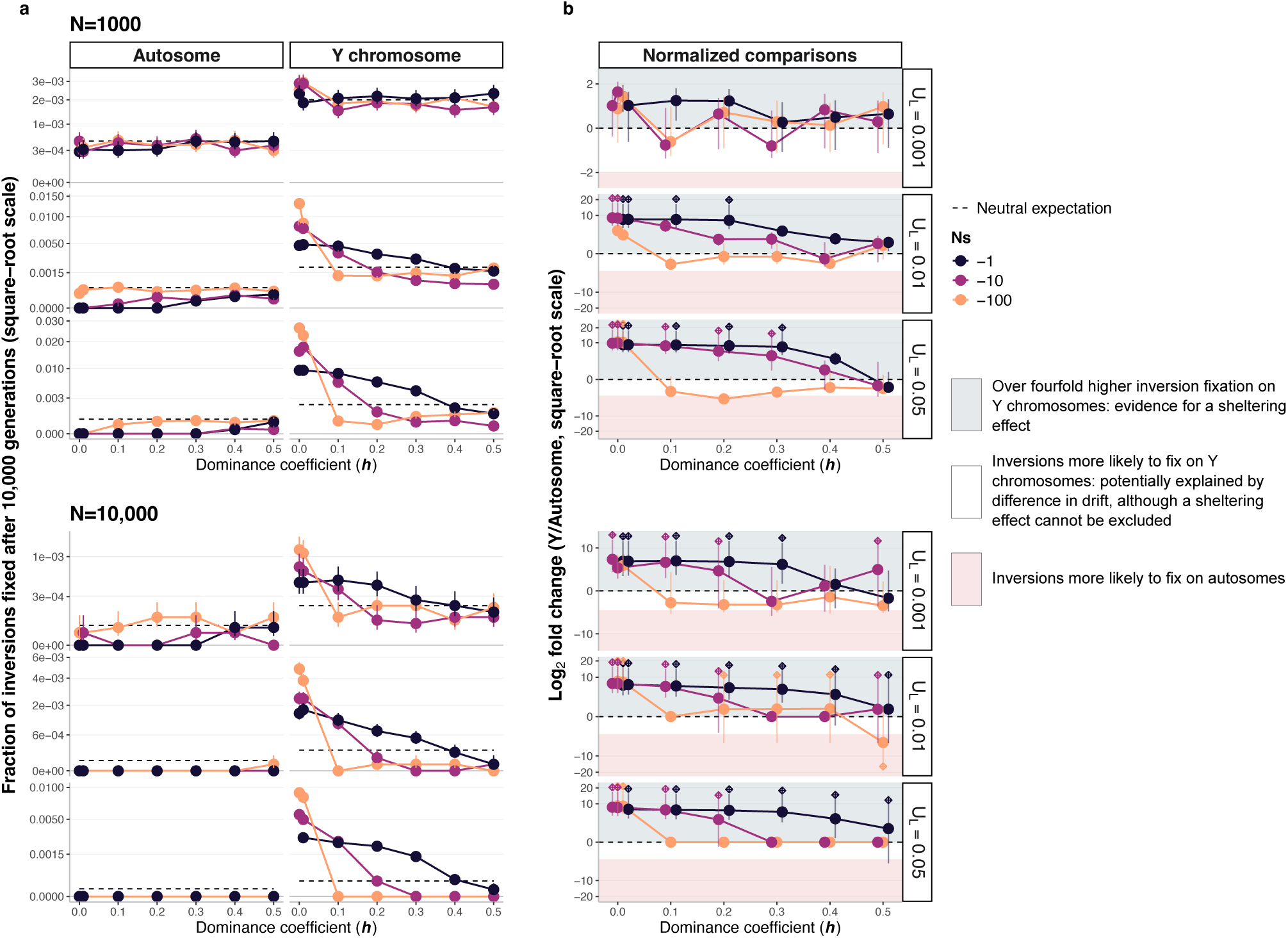
Less-loaded inversions are more likely to fix on Y chromosomes than on autosomes. **a.** Proportion of inversions that reached fixation after 10,000 generations in stochastic simulations with N = 1,000 and N = 10,000 across different combinations of parameter values (N, N*s, h, U_L_) after 6N generations. For each parameter combination, 50,000 inversions capturing a random genomic fragment were simulated. Dashed lines represent the neutral expectation, i.e. 2/N for Y chromosomes and 1/(2N) for autosomes. In contrast to Figure 3c in Jay et al. (2022), all inversions are shown here, rather than only the loaded inversions that survived the first 20 generations. Error bars represent 95% binomial confidence intervals; none are shown when no fixation occurred. **b.** Log_2_ ratio of the normalized fixation probabilities for Y-linked versus autosomal inversions. Absolute fixation probabilities (shown in panel a) were normalized by dividing them by their respective neutral expectations. Values above 0 indicate that inversions had more than fourfold higher absolute fixation probability on Y chromosomes (consistent with a sheltering effect), whereas values below −2 indicate higher absolute fixation probability on autosomes. To represent cases in which no autosomal or Y-linked inversion fixed in the simulations—which would otherwise produce an infinite ratio—we assumed that 0.5 inversions fixed in those cases. These situations are indicated by diamonds above or below the corresponding points. Diamonds above 0 indicate scenarios where no inversion fixed on autosomes, whereas diamonds below 0 indicate that no inversion fixed on the Y chromosome. Error bars show the 95% credible interval of the log2 fold change estimated from Monte Carlo sampling (see Methods). Figure S4 show identical analyses except that Y-linked inversions were simulated for 6N/4 generations instead of 6N generations, thereby accounting for differences in effective population size between autosomes and Y chromosomes.

**Figure 4.**
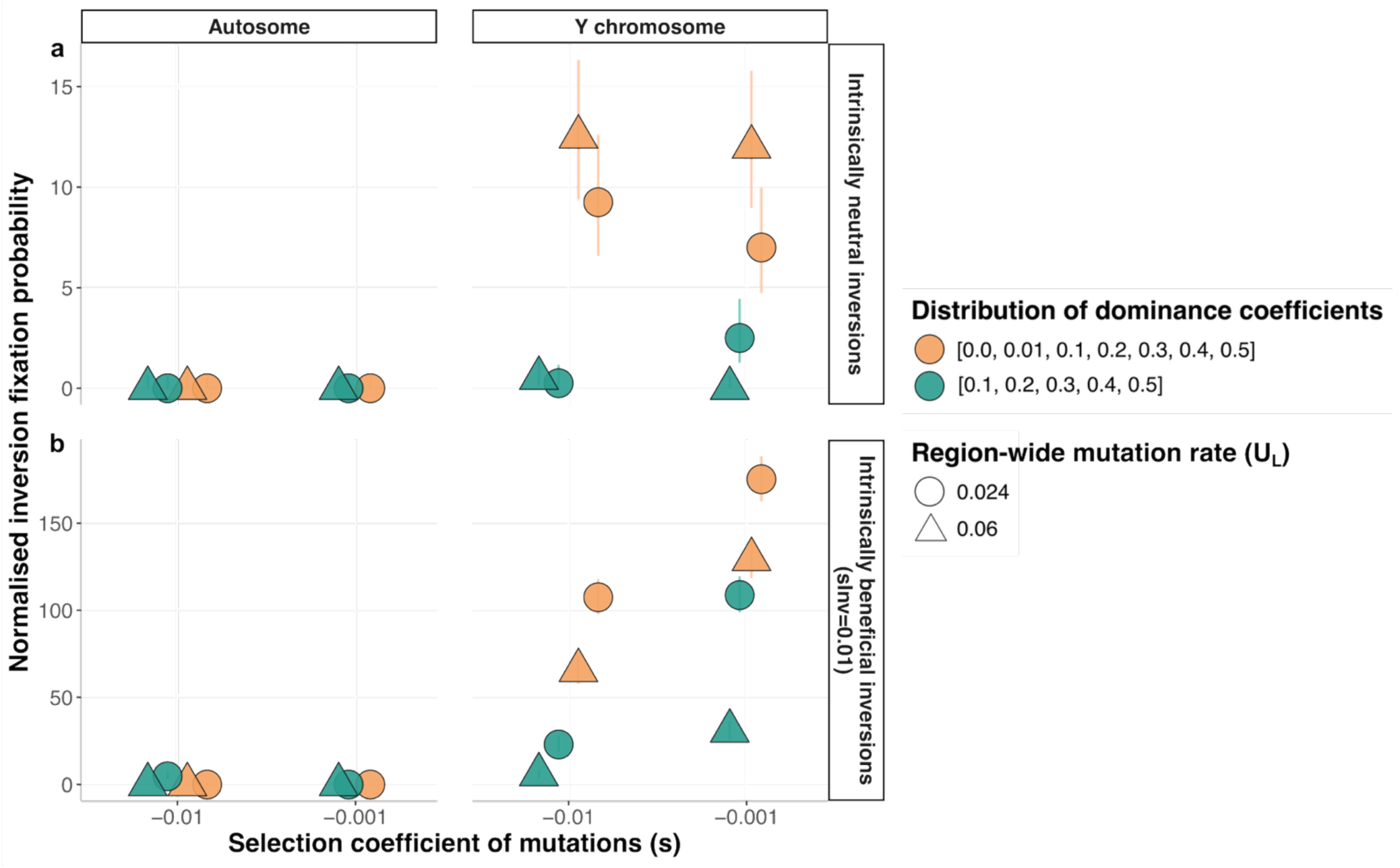
The sheltering effect facilitates inversion fixation on Y-like sex chromosomes when segregating mutations have a distribution of dominance coefficients and when inversions are intrinsically beneficial. **a,b,** Proportion of inversions that reached fixation after 6N generations in stochastic simulations divided by their respective neutral fixation probability (i.e., normalized probability), across different combinations of parameter values with N=12,500 or N=3,125. This figure reports simulations where mutations had their fitness coefficients sampled from a gamma distribution with mean −0.001 or −0.01 and shape 0.2, and their dominance coefficient randomly sampled with uniform probabilities among either [0.1, 0.2, 0.3, 0.4, 0.5] (mean=0.3, no fully recessive mutations), or [0.0, 0.01, 0.1, 0.2, 0.3, 0.4, 0.5] (mean=0.22, with fully recessive mutations). **b,** To show that the sheltering effect can act even in the absence of the lower-load advantage, we performed simulations of inversions benefiting from an intrinsic selective advantage (s_Inv_=0.01), in addition to the advantage or disadvantage conferred by their relative mutation load. A total of 100,000 inversions of 2Mb and 5Mb were studied for each combination of parameter values. In Figure S7, in addition to the simulations with N=12,500, we also performed simulations with an autosomal population of size equivalent to the effective population size of the Y chromosome, i.e. 3,125 (¼ of 12,500), as another approach to control for difference in population size between the Y chromosome and autosomes. Absolute fixation probabilities were normalized by dividing them by their respective neutral expectations, for controlling for population size (see Box 1a).

### Loaded inversions are much more likely to fix when they capture a Y-like sex-determining locus (Step 2)

Even though many inversions appear to carry a reduced mutational load, most inversions of moderate to large sizes still harbour some deleterious mutations—that is, they are rarely mutation-free (Figure 2c). The presence of these mutations may differentially influence the evolutionary trajectories of inversions on autosomes versus Y chromosomes. Indeed, inversions located on autosomes can be exposed in the homozygous form as they increase in frequency, whereas Y-linked inversions remain permanently heterozygous during their spread, thereby partially masking the deleterious effects associated with their mutation load, which corresponds to the sheltering effect. If mutation dynamics after the inversion formation are ignored, it is straightforward to show that less-loaded inversions tend to remain at low frequency when they occur on autosomes, whereas they can reach fixation on Y chromosomes due to this difference in heterozygosity (Supplementary text S1 and Figure S1).

Simulations of inversion frequency trajectories in finite populations with mutation accumulation following inversion formation confirmed the tendency identified in the deterministic model without mutation accumulation: over most of the parameter space explored, inversions were much more likely to fix if they captured the sex-determining allele on the Y chromosome than if they were located on autosomes (Figure 3a and Figures S2-3). On average, inversions were 22 times more likely to fix on Y chromosomes than on autosomes (Figures 3). Differences in inversion fixation probabilities between autosomes and the Y chromosome were more pronounced when segregating mutations were more recessive: fixation was for instance 52 times more likely on the Y chromosome than on autosomes when h≤0.1.

For a subset of parameter combinations, the probability of inversion fixation was higher than the expectation for a neutral variant (*i.e*., 1/2N for autosomes, 2/N for Y chromosomes; Figure 3). This was notably true on the Y chromosome when the segregating mutations were moderately to strongly recessive and weakly deleterious. This shows that the mere presence of deleterious mutations in genomes can change the fitness landscape in such a way that this facilitates inversion fixation. For the rest of the parameter combinations, the probability of inversion fixation on both the autosomes and the Y chromosome was lower than the expectation for a neutral variant (Figure 3). However, a fixation probability below the neutral expectation for a given parameter set does not, in any way, imply that the lower-load advantage or the sheltering effect do not contribute to inversion fixation on the Y chromosome under those conditions (see Box 1). It rather indicates that, under those conditions, the benefit associated with capturing fewer deleterious mutations was insufficient to offset other consequences of the occurrence of deleterious mutations in genomes, such as the progressive degeneration of non-recombining segments driven by drift. The presence of deleterious mutations can thus overall decrease inversion fixation probabilities compared to a situation without deleterious mutations. Nevertheless, even in these situations, the lower-load advantage and the sheltering effect can have important contribution to inversion dynamics.

To assess whether the permanent heterozygosity of the Y chromosome facilitates the fixation of loaded inversions through the sheltering effect, fixation probabilities on the Y chromosome must be compared to those on autosomes, while accounting for differences in effective population sizes between these genomic compartments (Box 1). Because the Y chromosome has one quarter of the effective population size of autosomes, differences in genetic drift alone can indeed generate differences in fixation probabilities even in the absence of any sheltering. To control for this effect, we normalized the inversion fixation probabilities by their neutral expectations for each chromosome type, *i.e*., 2/*N* for the Y chromosome and 1/2*N* for autosomes. A significantly higher normalized fixation probabilities on the Y chromosome than on autosomes provides evidence for a sheltering effect.

These normalized comparisons indicate that the sheltering effect contribute to inversion fixation when segregating mutations are partially recessive, with the magnitude of the effect increasing as dominance decreases (Figure 3). When mutations were strongly deleterious (*Ns* <= −100), only scenarios where mutations were highly recessive (*h* ≤ 0.01) produced higher normalized fixation probabilities on the Y chromosome relative to autosomes. In contrast, when segregating mutations were nearly neutral (*Ns* = −1), normalized fixation probabilities of inversions on the Y chromosome substantially exceeded those on autosomes even under moderately recessive mutations (*h* ≤ 0.5). Overall, we found evidence for a sheltering effect in 50% of the parameter space explored in Figure 3. Indeed, 73% of parameter sets exhibited higher normalized fixation probabilities on the Y chromosome than on the autosomes, and 50% showed non-overlapping confidence intervals.

Within genomes of natural populations, dominance coefficients vary among mutations. While several studies have estimated that the average dominance coefficient of deleterious mutations is around *h* ≈ 0.25 (Agrawal & Whitlock, 2012; Manna et al., 2011, 2012; Di & Lohmueller, 2024), the full distribution of dominance effects around this mean remains poorly characterized. To study how the dominance coefficient distribution affects the probability of inversion spread and fixation, we conducted simulations using two distributions, both with a mean close to *h* = 0.25, but one distribution allowing for fully recessive mutations, and the other excluding them (Figure 4a and S6). In both scenarios, we observed that inversions had much higher normalized fixation probabilities on the Y chromosome than on autosomes. This suggests that a subset of relatively recessive mutations (*h* ≤ 0.2) is sufficient to generate differences in fixation dynamics between Y-linked and autosomal inversions due to the sheltering effect.

To further emphasize that the sheltering effect (step 2) and the lower-load advantage (step 1) represent distinct selective effects, we performed additional simulations in which inversions, beyond their mutational load, carried an intrinsic selective advantage. This advantage could arise, for example, from gene disruption or altered gene expression caused by the inversion breakpoints (Dobzhansky, 1972; Kirkpatrick, 2010; Villoutreix et al., 2021) (Figure 4b and S6; see Methods). Taking the difference in drift between autosomes and the Y chromosome into account, we found that intrinsically beneficial inversions were more likely to fix on the Y-like sex chromosome in 100% of the parameter space explored in Figure 4 due to the sheltering of their recessive load.

As expected (Kimura & Ohta, 1969), we found that beneficial inversions took more generations to fix when population sizes were larger (Figure S6). This longer time favored the additional accumulation of deleterious mutations in inversions, decreasing their fitness and, in some cases, preventing their fixation (Figure S2-3). The probability of inversion fixation therefore decreased with increasing population size, both on autosomes and on Y chromosomes (Figures 3 and S2-3). However, this decrease was more pronounced in autosomes, resulting in a much larger difference in inversion fixation probabilities between the Y chromosome and autosomes when *N* = 10,000 (ratio 128:1 overall, 640:1 when *h*≤0.1) than when *N* = 1,000 (ratio 18:1 overall, 40:1 when *h*≤0.1; Figures 3a and S4). Note that, depending on parameter values, the lower fixation probabilities of inversions in larger populations may be offset by the higher number of inversions expected to arise in such populations.

### Other systems with permanently heterozygous alleles, other recombination modifiers and sex chromosome-autosome fusion

Further simulations showed that the sheltering effect facilitated the fixation of inversions under a broader range of conditions and around other types of loci under balancing selection. This was observed when: (i) two or more permanently heterozygous alleles segregated at a mating incompatibility locus, such as in plant self-incompatibility or fungal mating-type systems [Figure S8; (Hartmann et al., 2021; Takayama & Isogai, 2005)]; (ii) alleles were not permanently heterozygous but exhibited strong overdominance, as described for several supergenes [Figure S9; (Jay et al., 2021; Küpper et al., 2016; Wang et al., 2013)]; (iii) the fitness effects of mutations across the genome followed a gamma distribution, so that segregating mutations differed in their selective effects (Figure 4 and S10); (iv) recombination modifiers suppressed recombination even in the homozygous state, as is the case for epigenetic mechanisms such as histone modifications or DNA methylation (Boideau et al., 2021), rather than only in heterozygotes as for chromosomal inversions (Figure S8); and (v) species exhibited a haplodiplontic life cycle, with alternation of haploid and diploid phases (Figure S11).

In addition, our simulations indicate that the sheltering of partially recessive deleterious mutations can promote the fusion between a permanently heterozygous sex chromosome and an autosome. This occurs when chromosome fusions are associated with an extension of the non-recombining region onto the newly fused autosomal segment (Figure S12).

### Evolution of non-recombining sex chromosomes with evolutionary strata despite possible reversions (step 1 + step 2 + step 3)

We have shown that, due to the sheltering effects, inversions are often much more likely to fix when they capture a permanently heterozygous allele than when occurring elsewhere in the genome. This process should thus lead to the repeated fixation of inversions around permanently heterozygous alleles, such as the male-determining allele in XY systems, leading to the formation of non-recombining sex chromosomes with a typical pattern of evolutionary strata. We investigated the formation of evolutionary strata through the successive fixation of multiple, potentially overlapping inversions, by simulating the evolution of large chromosomes experiencing recurrent inversion events, using parameter values representative of those observed in mammals (Figures 5, 6 and S13). A key focus was the potential for recombination restoration via inversion reversion, which may be favored when a fixed inversion on the Y chromosome accumulates a sufficiently high mutational load to exceed the population average (Lenormand & Roze, 2022).

**Figure 5.**
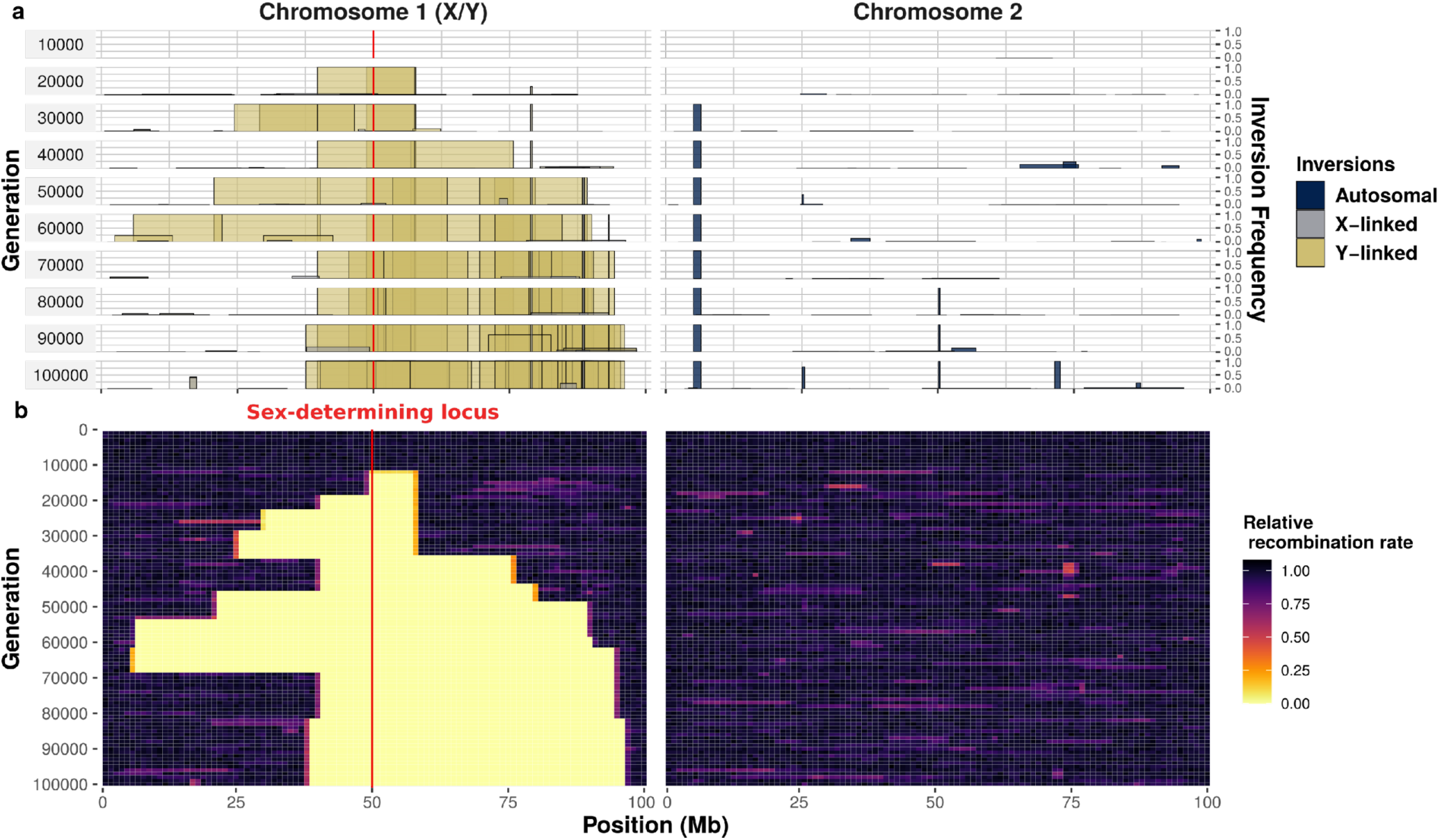
Accumulation of successive inversions around a Y-like sex-determining allele in an XY system, leading to the formation of non-recombining sex chromosomes with evolutionary strata. Results of a simulation of N=1000 individuals, each with two pairs of 100 Mb chromosomes, over 100,000 generations. Chromosome 1 harbors an X/Y sex-determining locus at 50 Mb (individuals are XX or XY). In each generation, one inversion appears, on average, in the whole population, in an individual sampled uniformly at random, with the two recombination breakpoints sampled uniformly at random from k=100 potential breakpoints. **a.** Overview of chromosomal inversion frequency and position for 10 different generations. The width of the box represents the position of the inversion and the height of the box indicates inversion frequency. Inversions appearing on the Y chromosome are depicted in yellow, those appearing on the X chromosomes are depicted in gray. The colors are not entirely opaque, so that regions with overlapping inversions appear darker. Previously fixed inversions may be lost due to the occurrence of beneficial reversions and selection. **b.** Changes in the relative rate of recombination over the entire course of the simulation. The numbers of recombination events occurring at each position (binned in 1 Mb windows) are recorded at the formation of each offspring, across all homologous chromosomes in the population. Only recombination events between the X and the Y chromosomes are shown for chromosome 1 (i.e., recombination events between the two X chromosomes in females are not shown). Unlike chromosome 1, chromosome 2 harbors no permanently heterozygous alleles. All inversions on this chromosome suffer from homozygote disadvantage and very few inversions therefore become fixed on chromosome 2. See Figure S13 for a simulation with N=10,000 individuals. The datasets and scripts used to produce this figure are available on Figshare (doi:10.6084/m9.figshare.19704457) and GitHub.

**Figure 6.**
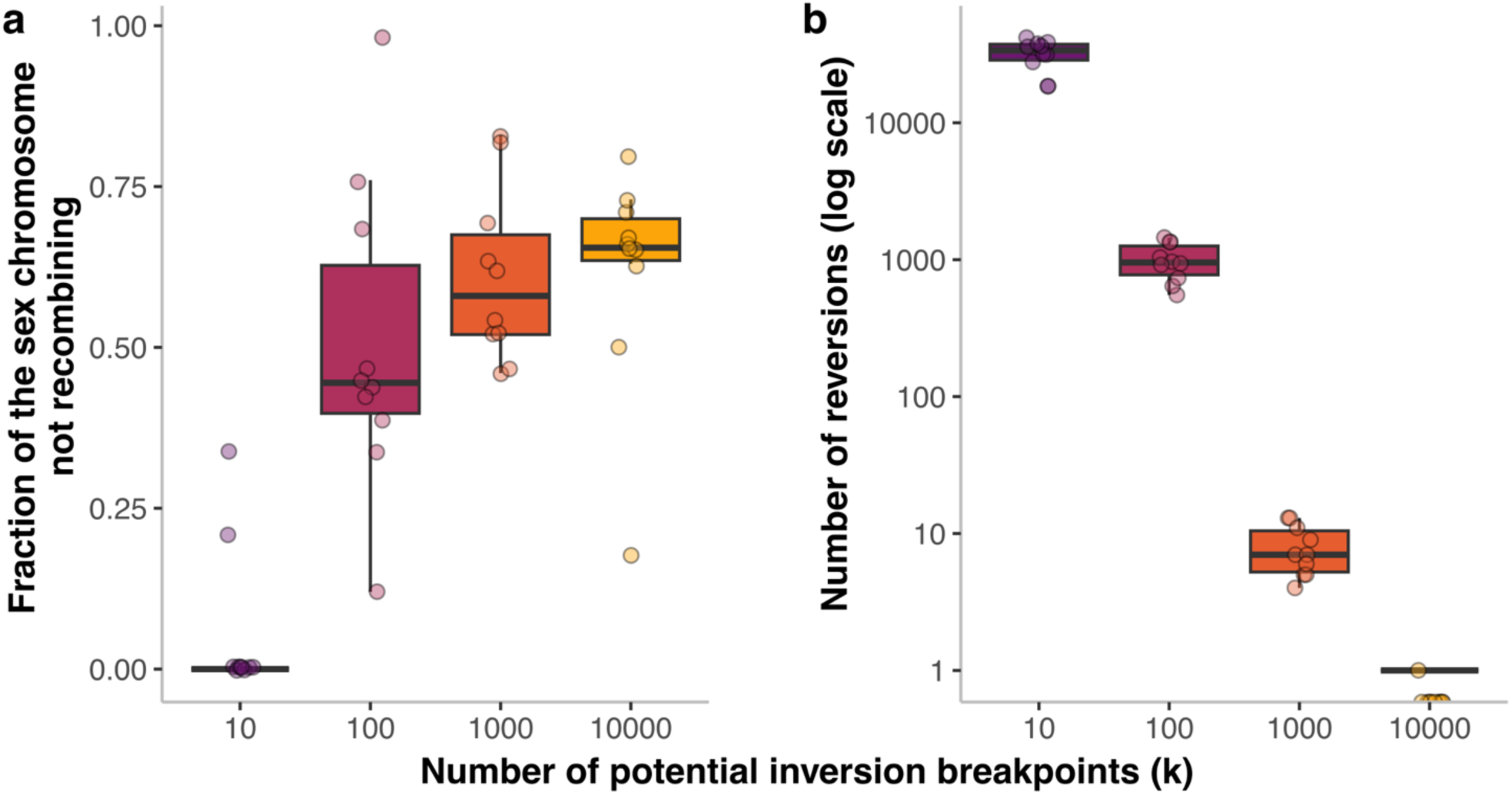
Effect of the number of potential inversion breakpoints on the evolution of recombination suppression in sex chromosomes. Each dot represents the result of a simulation with N=1000 individuals. For each number of breakpoints, 10 simulations were conducted. See Figure 5 for an example of such a simulation. **a.** Fraction of the length of the Y sex chromosome not recombining after 100,000 generations. **b.** Number of reversions occurring over the course of the 100,000 generations. Boxplot elements: central line: median, box limits: 25th and 75th percentiles, whiskers: 1.5x interquartile range. The dataset and script used to produce this figure are available on Figshare (doi:10.6084/m9.figshare.19704457) and GitHub.

We simulated, over 100,000 generations, populations of *N*=10,000 or *N*=1,000 individuals, carrying two 100 Mb chromosomes, one of which harbored a mammalian-type sex-determining locus (XY males and XX females). Individuals experienced only deleterious or weakly deleterious mutations, with mutation rates and fitness effects similar to those observed in humans (fitness effect being drawn from a gamma distribution). The dominance coefficient of each mutation was chosen uniformly at random from a wide set of values (see Methods for details). At the start of each simulation, we randomly sampled *k* genomic positions that could be used as inversion breakpoints, with *k* being 10, 100, 1000 or 10,000. These genomic positions represent inversion hotspots, such as those that can be generated by repeats in genomes (Porubsky et al., 2022). In each generation, we introduced *j* inversions, *j* being sampled from a Poisson distribution. To limit simulation times, we used high inversion rates, with one inversion occurring on average each generation in the whole population. The two breakpoints of each inversion were chosen at random from the *k* positions. It was, therefore, possible for two independent inversions to appear at the same position, allowing, in particular, the reversion of an inversion to its ancestral orientation, thereby restoring recombination. We assumed that inversions partially overlapped by another inversion or captured by a larger inversion could not be reversed, *i.e*., that recombination could not be restored in such situations even if subsequent inversions re-used the same breakpoints. Indeed, reversions of partially overlapped inversions do not restore ancestral arrangements but instead result in complex reshufflings of gene order and orientation and are therefore unlikely to restore recombination. We ran 10 simulations for each set of parameters.

In all simulations assuming relatively large numbers of potential inversion breakpoints (*k*=100, 1000, 10,000), the Y chromosome progressively stopped recombining with the X chromosome as it accumulated successive inversions fully linked to the male sex-determining allele (Figures 5, 6 and S13). After the occurrence of an initial inversion capturing the Y sex-determining allele, multiple inversions partially overlapping this inversion or other Y-fixed inversions were selected for, thereby generating a growing chaos of overlapping chromosomal rearrangements. The non-recombining region, thus, extended around the sex-determining locus in a stepwise manner, perfectly reflecting the evolution of sex chromosomes and other supergenes with evolutionary strata (Figure 5). Some events in the gradual extension of recombination suppression were reversed, due to the occasional occurrence of beneficial reversions (Figures 5 and 6). The accumulation of overlapping inversions was, however, more rapid than the occurrence of beneficial reversions, leading to a progressive extension of the non-recombining region (Figures 5 and 6). By contrast, when we assumed a smaller number of potential breakpoints (*k* = 10), recombination suppression between sex chromosomes evolved in only two of the 10 simulations, and over only a small genomic region (Figure 6), owing to the more frequent occurrence of beneficial reversions.

## Discussion

### Evolution of recombination suppression around permanently heterozygous alleles through the combination of a lower-load advantage and the sheltering effect

Our results show that recombination suppression can evolve on sex chromosomes and other supergenes, because of three phenomena linked to the widespread presence of partially recessive, deleterious variants in genomes: i) inversions (or any recombination suppressor) can be favored solely because they contain fewer deleterious mutations than the population average, a situation applying to a substantial fraction of the inversions formed; ii) such inversions tend to display overdominance: they are beneficial in the heterozygous state but suffer from a homozygote disadvantage, which prevents them from reaching high frequencies and becoming fixed on autosomes; iii) when, by chance, such less-loaded inversions capture a permanently heterozygous allele, they do not suffer from this homozygote disadvantage and are therefore able to increase in frequency until they are fully associated with the permanently heterozygous allele (*e.g*., they become fixed in the Y chromosome population). These three phenomena have been reported independently in several studies, but, to our knowledge, never in interaction [see (Lenormand & Roze, 2022; Nei et al., 1967; Olito & Abbott, 2024) for i), (Kirkpatrick, 2010; Ohta, 1971) for ii), and (Antonovics & Abrams, 2004; Hartmann et al., 2021) for iii)]. The combined influence of the mechanisms related to ii) and iii) has been shown to promote sex chromosome-autosome fusion in highly inbred populations (Charlesworth & Wall, 1999). We show here that such mechanisms can readily lead to the stepwise extension of the non-recombining region on sex chromosomes themselves, without the need for inbreeding. Moreover, unlike previous studies [*e.g*. (Charlesworth et al., 1987; Connallon et al., 2018; Olito & Abbott, 2024)], we show that the higher probability of inversion fixation on Y chromosomes is not restricted to mutation-free inversions, but applies to any inversion loaded with deleterious recessive mutations.

The theory proposed here to explain the stepwise evolution of recombination suppression applies to any locus with at least one permanently or nearly permanently heterozygous allele. It only requires that, within a population, individuals carry different numbers of partially recessive deleterious mutations in a genomic region that can be subjected to recombination suppression. As discussed below, this situation is probably frequent in diploid, dikaryotic or heterokaryotic organisms (Berdan et al., 2023; Giner-Delgado et al., 2019; Porubsky et al., 2022; Wellenreuther & Bernatchez, 2018).

It is important to emphasize that the lower-load advantage and the sheltering effect are two distinct and independent mechanisms that can influence the evolutionary fate of inversions. The lower load corresponds to a selective advantage, arising when an inversion captures by chance a genomic segment carrying fewer deleterious mutations than the population average in this genomic region. The fitness benefit conferred by such an inversion increases with the absolute difference in mutation load between the captured segment and the population mean. Consequently, larger inversions—by capturing more sites—have a wider fitness distribution, and can thus experience a stronger selective advantage.

The sheltering effect corresponds to the protection from the homozygote disadvantage incurred by inversions loaded with partially recessive deleterious mutations when they become frequent. Indeed, autosomal inversions carrying partially recessive deleterious mutations experience a reduction in fitness as their frequency increases, as they increasingly occur at the homozygous state. In contrast, inversions linked to the permanently heterozygous Y chromosome are protected from this homozygous disadvantage, allowing even highly loaded inversions to spread and potentially fix. As shown in our analyses, this difference in fate between autosomal and Y-linked inversions due to the sheltering effect does not only apply to less-loaded inversions. It applies to any inversion, regardless of whether its spread is driven by genetic drift or a selective advantage, as long as it carries recessive deleterious mutations. Consequently, this effect is expected to facilitate the spread of virtually any inversion of substantial size on Y-like chromosomes.

### The right controls to study the sheltering effect

In this manuscript, as well as in Jay et al. (2022), we evaluated the efficacy of the sheltering effect by comparing the fate of inversions with deleterious recessive mutations on the Y chromosome to their fate on autosomes, and not by a comparison with situations where all mutations are neutral (*s*=0). Indeed, recessive deleterious mutations prevent the fixation of most inversions on autosomes (an effect which is absent when *s*=0), while the sheltering effect protects inversions from this fitness loss on Y-like chromosomes. The sheltering effect thus saves inversions specifically on Y-like chromosomes, when deleterious mutations are segregating. We therefore need deleterious mutations to be present to be able to evaluate to what extent their sheltering contributes to inversion fixation (see Box 1; see also Box 3 in Jay et al., 2024, and discussion in Saunders & Muyle, 2024). The comparison between autosomes and the Y chromosome, controlling for their different population effective sizes, allows us to assess whether the sheltering effect can be a significant contributor to inversion fixation on Y-like chromosomes in populations when recessive deleterious mutations are present. Because deleterious mutations are known to segregate in natural populations (Agrawal & Whitlock, 2012; Eyre-Walker & Keightley, 2007), these effects need to be considered if we are to understand the evolution of recombination suppression on sex chromosomes in nature.

### Conditions for the occurrence of the lower-load advantage and the sheltering effect

In this study, we explored a broad parameter space, with *N* ranging from 1,000 to 12,500; *s* from −0.0001 to −0.1; *h* from 0 to 0.5; *μ* from 1e-09 to 1e-08; and inversion sizes from 500kb to 5 Mb (or region-wide mutation rate *U_L_* from 0.0005 to 0.05). Our results demonstrate that the number of segregating deleterious mutations and their dominance coefficients are crucial factors for the occurrence of the lower-load advantage and the sheltering effect, with more mutations and more recessive mutations favoring inversion fixation.

Numerous studies have shown that a substantial proportion of new and segregating mutations are deleterious (Agrawal & Whitlock, 2012; Eyre-Walker & Keightley, 2007). While the precise numbers and effects of these mutations are debated, it is widely accepted that genomes carry tens of thousands of harmful mutations. Therefore, any large inversion (*e.g*., 1 Mb) is expected to harbor multiple deleterious mutations. For instance, the average human genome contains approximately 4.1 to 5.0 million polymorphic sites, with an estimated 25% of these mutations being deleterious (Racimo & Schraiber, 2014). Even if this estimate was overestimated by a factor of 100, megabase-scale inversions would still contain multiple deleterious variants, setting the stage for the lower-load advantage and the sheltering effect. Importantly, megabase-scale inversions, as we modelled, are commonly observed in natural populations. For instance, the two most recent evolutionary strata on the human Y chromosome span approximately 1 and 4 Mb, respectively (Zhou et al., 2023), which corresponds to *U_L_*=0.0007 and *U_L_*=0.0028 when considering a human genome-wide deleterious mutation rate of 2.2 (Agrawal & Whitlock, 2012).

Although empirical estimates of the dominance coefficient for mutations in natural populations are limited, studies in *Drosophila*, yeasts and nematodes have estimated the average dominance coefficient for deleterious mutations at approximately *h* = 0.25 (Agrawal & Whitlock, 2012; Manna et al., 2011; Di & Lohmueller, 2024). This value may seem to lie near the boundary of the conditions under which the sheltering effect is expected to operate (depending on the distribution of selective coefficients). However, it is not the average dominance coefficient that matters for the sheltering effect, but instead the proportion of moderately to strongly recessive mutations, that is, the full distribution of dominance coefficients. Inversions arising in taxa with a substantial proportion of strongly recessive mutations are expected to experience a strong homozygous disadvantage on autosomes, which in turn translates into a strong sheltering effect on the Y chromosome, even if the average dominance coefficient is high. While the distribution of dominance coefficients in nature is poorly documented, an average dominance coefficient for deleterious mutations at *h* = 0.25 suggests that many deleterious mutations have dominance coefficients well below 0.25. For example, yeast gene knockout data led to estimates of average dominance coefficient of 0.046 for mutations affecting catalytic functions (Agrawal & Whitlock, 2011). There is also substantial evidence that the distribution of dominance coefficients for strongly deleterious mutations is right-skewed, with an overrepresentation of strongly recessive mutations (Mrnjavac et al., 2025; Di & Lohmueller, 2024). Several studies that jointly estimated *h* and *s* have shown that more strongly deleterious mutations generally tended to be more recessive (Di & Lohmueller, 2024).

Such observations suggest that the conditions required for the sheltering effect to operate could be regularly met in natural populations. In particular, we show that the sheltering effect can operate when inversions contain many weakly deleterious mutations (*Ns* = −1), even when these mutations are only moderately recessive (*h* ≤ ∼0.4). In contrast, when genomes carry only strongly deleterious mutations (*e.g*., *Ns* = −100), the sheltering effect is observed only when these mutations are more strongly recessive (*e.g*., *h* ≤ 0.1). Therefore, if genomes harbor a large number of moderately recessive, weakly deleterious mutations, together with any proportion of more strongly deleterious and more recessive mutations, the sheltering effect could have a strong impact on inversion fates. However, additional empirical data are needed on the distribution of dominance and selective coefficients in natural populations, as well as on their variation across taxa, genes, and mutation types, to assess whether the conditions under which the sheltering effect substantially influences inversion dynamics are met.

### The rate of inversion fixation and the tempo of recombination suppression

Evolutionary strata rarely arise in natural populations. For example, the mammalian Y chromosome has experienced only five successive events of recombination suppression over approximately 250 million years (Cortez et al., 2014; Zhou et al., 2023). Consequently, models aiming to explain the evolution of sex chromosomes do not need to generate high rates of inversion fixation (i.e. of strata evolution), either in absolute terms or relative to neutral expectations.

Recent studies suggest that inversions may not be exceedingly rare and that their breakpoints are widely distributed across genomes (Hämälä et al., 2021; Porubsky et al., 2022; Todesco et al., 2020; Zhou et al., 2019). However, because empirical estimates of inversion occurrence rates in natural populations remain scarce, it is currently difficult to assess whether the fixation probabilities observed in our simulations are consistent with inversion fixation rates observed in natural population. Additional data on inversion occurrence rates are therefore needed to evaluate this aspect. In our simulations of the stepwise evolution of recombination suppression (Figures 5 and 6), we used relatively high inversion rates to make the computations tractable and to allow the stepwise extension of non-recombining regions to be observed within 100,000 generations. While different inversion rates would lead to shorter or longer timescales for the extension of non-recombining regions, they are not expected to qualitatively alter the outcome

### Mutation accumulation does not necessarily prevent recombination suppression extensions

On autosomes, we show that inversions maintained at low frequencies due to their homozygous disadvantage are typically rapidly lost, as they continue to accumulate additional deleterious mutations over time. On the Y chromosome, however, the rate of mutation accumulation following the formation of an inversion can be slow enough that it does not fully offset the initial selective advantage of less-loaded inversions. This can allow Y-linked inversions to reach fixation, despite ongoing degeneration. However, due to computational limitations, our analyses were restricted to relatively small population sizes (up to N=12,500). In larger populations, the time to fixation may be considerably longer, potentially allowing more deleterious mutations to accumulate before fixation occurs—thereby reducing the likelihood that such inversions will ultimately fix. Yet, our analyses show that the difference in normalized inversion fixation probabilities between the Y chromosome and autosomes increases with population size, suggesting that the lower-load advantage and the sheltering effect become more important relative to drift in shaping inversion fixation dynamics as population size increases. Recent analytical results (Offenstadt et al. 2026) support this pattern, showing that overdominant loci (such as less-loaded inversions) are more likely to fix over realistic timescales on Y chromosomes than on autosomes, and that this difference in fixation probability increases with population size. However, to more precisely understand the evolutionary dynamics of inversions in large populations, analytical models that explicitly incorporate the complex patterns of mutation accumulation—particularly the effects of drift and selective interference among mutations—are required, as the individual-based simulations used here become computationally intractable at larger population sizes.

### Long-term persistence of recombination suppression on sex chromosomes

Our results show that deleterious mutations accumulate following Y-linked inversion fixation, as observed on many non-recombining sex chromosomes (Bachtrog, 2013). This may lead to selection for inversion reversion, thereby restoring recombination (Lenormand & Roze, 2022). The question as to whether this actually occurs is common to all mechanisms explaining evolutionary strata, including sexual antagonism.

In this paper we showed that overlapping genomic rearrangements accumulating following recombination suppression could prevent recombination to be restored, depending on the number of putative inversion breakpoints. Indeed, partially overlapping inversions result in a complex reshuffling of gene order and orientation, preventing the restoration of recombination even in situations in which reversion could be selected for. When the number of potential breakpoints is relatively high, the probability of partially overlapping inversions occurring is higher than the probability of a reversion occurring, regardless of the relative rates of inversions and deleterious mutations.

The reversion of inversions has in fact been reported only in rare cases in which inversions occur at specific breakpoints rich in repeated elements (Cui et al., 2012; Hanson et al., 2014), as in our simulations when assuming only a few possible breakpoints in the genome. The number of genomic positions at which inversions can occur in natural conditions is unknown, but this number is likely to be high given the chaos of rearrangements observed in some sex and mating-type chromosomes with hundreds of different breakpoints (Badouin et al., 2015; Branco et al., 2017; Carey et al., 2021; Porubsky et al., 2022). A recent study reported the recurrent appearance of inversions at the same positions, but also the existence of numerous potential breakpoints of inversions in human genomes (Porubsky et al., 2022). Moreover, several studies have shown that chromosomal rearrangements rapidly accumulate in recently established regions of non-recombination (Badouin et al., 2015; Carpentier et al., 2022) and overlapping inversions are observed on many sex chromosomes, which should prevent the restoration of recombination by reversions (Bellott et al., 2014; Branco et al., 2017; Carey et al., 2021; Lemaitre et al., 2009; Skinner et al., 2021). Therefore, over a wide range of realistic parameter values, the reversion of inversions should not prevent the stepwise formation of non-recombining sex chromosomes. However, additional data on inversion frequencies in non-recombining regions, as well as on the rate at which reversions occur across a broad range of organisms, are needed to determine whether the accumulation of overlapping rearrangements is sufficiently rapid to prevent the restoration of recombination.

It should be noted that the lower-load advantage and the sheltering effect are expected to operate similarly for any cis-acting recombination suppressor, not only chromosomal inversions. However, other mechanisms known to affect recombination rates, such as epigenetic modifications, may be more readily reversible than chromosomal inversions and are unlikely to generate chaotic structures preventing the restoration of recombination. Consequently, the stability of recombination suppression may vary depending on the underlying mechanism. Some young non-recombining regions on mating-type chromosomes appear to be collinear, showing that they were not formed by inversions (Branco et al., 2018; Sun et al., 2017; Vittorelli et al., 2023; De Filippo et al., 2026), but further data on the causes of recombination suppression in sex chromosomes and other supergenes are needed to better understand their short- and long-term evolutionary dynamics.

Lenormand & Roze, 2024, argued that, in a longer term and without dosage compensation, the fitness of individuals carrying the inversions having accumulated a further load should decrease to a point where species could go extinct, and therefore that the combined effect of the lower-load advantage and sheltering effect could not explain the evolution of recombination suppression on sex chromosomes. However, that recombination suppression on sex chromosomes can lead to species extinction or sex chromosome turn-over unless dosage compensation or other mitigation mechanisms eventually evolve does not in itself negates the potential role of the less-loaded advantage, of the sheltering effect and of overlapping rearrangements in allowing the evolution and medium-term maintenance of recombination suppression (see also the discussion in Saunders & Muyle, 2024). Conversely, our theory does not negate a possible role of dosage compensation for the long-term maintenance of sex chromosomes, but it shows that *early* dosage compensation or sexual antagonism may not be required to explain sex chromosome evolution, in contrast to the conclusions reached in previous studies (Lenormand & Roze, 2022, 2024). A similar case may be asexuality, which can be selected for in the short term but can lead to species extinction in the long term (de Vienne et al., 2013). Furthermore, the early dosage compensation hypothesis requires a form of sexual antagonism, as it assumes the existence of numerous sex-specific gene regulators for genes on proto-Y sex chromosome, even before the evolution of recombination suppression (Jay et al., 2024; Lenormand & Roze, 2022). This early dosage compensation mechanism (Lenormand & Roze, 2022) cannot, therefore, explain evolutionary strata on fungal mating-type chromosomes, with which no form of antagonistic selection is associated (Bazzicalupo et al., 2019; Hartmann et al., 2021). Further investigation on the existence of dosage compensation in very young evolutionary strata is needed to assess whether early dosage compensation is required for their evolution. Studies in plants, birds and reptiles, for example, have actually suggested that young evolutionary strata often do not display dosage compensation (Jay et al., 2024; Sacchi et al., 2025).

### Predictions regarding the variability of sex-chromosome structure across species

We show that recombination suppressors such as inversions are more likely to spread and fix in regions harboring many segregating deleterious mutations, in which they have a greater chance of capturing highly advantageous haplotypes. Our model therefore predicts that species harboring a large number of deleterious recessive variants, due to their small population size, short haploid phase, outcrossing mating system, high mutation rate and high levels of mutation recessiveness, for example, will be more prone to the evolution of large non-recombining regions with evolutionary strata on sex chromosomes than species with a low mutation load. In addition, in species with large population sizes, the time required for an inversion to become fixed may exceed the time during which the number of deleterious mutations in the inversion remains below average. This should prevent some inversions from becoming fixed, potentially decreasing the expansion rate of non-recombining regions on sex chromosomes in species with large population sizes. Depending on mutation effects and on the relative rates of inversions and mutations, this could however be compensated by the occurrence of a higher number of inversions each generation in large populations.

Consequently, variations in population size, mutation rate, length of haploid phase and mating system (outcrossing versus selfing, or inbreeding) across lineages may account for the large variation in sex-chromosome structures in nature, with some organisms maintaining homomorphic sex chromosomes and others evolving highly differentiated sex chromosomes with multiple evolutionary strata (Abbott et al., 2017).

The more efficient purging of recessive deleterious mutations in species with an extended haploid phase (Kondrashov & Crow, 1991; Scott & Rescan, 2017), and the more limited sheltering possibility in such lifecycles, could therefore potentially account for the smaller non-recombining regions observed on the sex chromosomes of plants and algae than in animals (Coelho et al., 2018; Filatov, 2015). In fungi for example, multiple species with a diploid-like life cycle have repeatedly and independently evolved stepwise recombination suppression around mating-type loci in diverse lineages, but not their closely related species with haploid-like life cycles (De Filippo et al., 2026; Hartmann et al., 2021; Jay et al., 2024).

A test of our model could thus look for association between the size of non-recombining regions or the number of evolutionary strata on sex chromosomes and estimates of the number of deleterious mutations segregating in genomes, or parameters predicting the accumulation of deleterious mutations, such as population size, mating systems, length of the haploid phase and mutation rates (Jay et al., 2024).

## Conclusion

Our model, based on the observation that partially recessive deleterious mutations are widespread in genomes, can contribute to explain the evolution of stepwise recombination suppression on sex chromosomes. Furthermore, it can explain why some supergenes, such as fungal mating-type chromosomes and autosomal supergenes in butterflies and ants, display evolutionary strata (Carpentier et al., 2022; De-Kayne et al., 2025; Jay et al., 2021; Yan et al., 2020; Branco et al., 2017; Hartmann et al., 2021). In addition, our framework may explain why meiotic drivers, which are often permanently heterozygous, are frequently associated with extended non-recombining regions involving polymorphic chromosomal inversions (Dyer et al., 2007; Reinhardt et al., 2014).

Our model therefore provides a general and simple framework that can contribute to understanding the evolution of non-recombining regions around a wide range of loci carrying permanently or near-permanently heterozygous alleles. The different theoretical frameworks proposed to explain the evolution of sex chromosomes, as reviewed in Jay et al. (2024), are not mutually exclusive, and their relative contributions likely depend on underlying population genetic parameters. Further empirical work is therefore needed to better characterize these parameters, such as the distribution of dominance and selection coefficients and the frequency of sex-antagonistic mutations in natural populations. Linking the presence and number of evolutionary strata observed across taxa to these parameters will provide a direct and testable way to evaluate the relevance of our model and alternative hypotheses.

## Materials and Methods

### Simulation of inversion trajectory on autosomes and Y chromosomes

We used SLiM V4.3 (Haller & Messer, 2019) to simulate the evolution of a single panmictic population of *N* individuals in a Wright-Fisher model, *N* being either 1000 or 10000. To assess the fate of inversions under various conditions (Figure 3), we simulated individuals with one pair of 5Mb chromosomes on which mutations occurred at a rate *μ*, with *μ* being either 10^-8^ or 10^-9^ per bp, their dominance coefficient *h* ranged from 0 to 0.5 (0, 0.01, 0.1, 0.2, 0.3, 0.4, 0.5) and their selection coefficient *s* from −0.1 to −0.0001, so that *Ns* was equal either to −1, −10 or −100 (note that scenarios with *N*=1000 and *s*=-0.0001, resulting in Ns=-0.1, were not considered in the analyses since the mutations then behaved as if they were neutral in these cases). We considered a recombination rate of 10^-7^ per bp.

For each parameter combination (*u*, *h*, *s, N*), a simulation was run for 6*N* generations, to allow the population to reach an equilibrium for the number of segregating mutations (Figure S14). The population state was saved at the end of this initialization phase. These saved states (one for each parameter combination) were repeatedly used as initial states for studying the dynamics of recombination modifiers. Several cases were then considered. For the autosomal cases, nothing additional was done. For others cases, a single locus subject to balancing selection was introduced at the chromosome center, for which two situations were considered: (i) the locus had two alleles, only one of which was permanently heterozygous, mimicking a classical XY (or ZW) determining system (Figure 3, 4, S4-7,10-12), (ii) the locus had two permanently heterozygous alleles, mimicking, for instance, the situation encountered at most fungal mating-type loci (Figure S8).

Recombination modifiers mimicking inversions of size *n*=1Mb and *n*=5Mb were then introduced, centred on the chromosome (and therefore around the locus under balancing selection in the non-autosomal cases), encompassing a region with region-wide mutation rate *U_L_*=*n*\**μ* which was therefore equal to 0.001, 0.01 or 0.05 (scenarios with *U_L_*=0.005 were not analysed to save computation time). For each parameter combination (*h*, *s*, *u*, *N*, heterozygosity rule, size of the region affected by the recombination modification and position on the genome), we ran 50,000 independent simulations starting with the introduction of a single recombination modifier in the same saved initial population. These inversion-mimicking, recombination modifier mutations were introduced on a single, randomly selected chromosome and, when heterozygous, they suppressed recombination across the region in which they resided (*i.e*., as a *cis*-recombination modifier).

We monitored the frequency of these inversion-mimicking mutations during 6*N* generations, during which all evolutionary processes (such as point mutation, recombination and mating) remained unchanged, *e.g*., mutations were still appearing on inversions following their formation. In order to reduce the simulation time, simulations were stopped when the inversions reached fixation, *i.e*., when the inversion reached a frequency of 1.0 on autosomes or 0.25 on sex chromosomes. Under the same assumptions, but for a reduced set of combinations of parameter values, we also studied the dynamics of recombination modifiers suppressing recombination also when homozygous and not only when heterozygous, again across a fragment in which they resided (Figure S8).

One may argue that, because the effective population sizes of sex chromosomes and autosomes differ, using the same simulation duration for Y chromosomes and autosomes could bias comparisons of inversion fixation probabilities in favor of Y-linked inversions. Whether this constitutes a bias, however, depends on the question being addressed—namely, whether one is interested in fixation probabilities over a given time frame or in their asymptotic values (i.e., after an infinite number of generations). We therefore additionally present in figure S4 results obtained after 6N generations on autosomes but only 6N/4 generations on the Y chromosome, thereby accounting for differences in effective population size between these genomic compartments. As shown by comparing figure S4 and figure 3, the resulting patterns are qualitatively unchanged, indicating that using 6N rather than 6N/4 generations for Y-linked inversions does not affect our conclusions.

### Simulations with a distribution of selective and dominance coefficients

We also simulated populations (Figure 4, S7 and S10) where each new mutation had its selection coefficient *s* drawn from a gamma distribution with a shape of 0.2 and a mean of −0.1, −0.01 or −0.001; its dominance coefficient of mutation (*h)* was drawn randomly with a uniform probability either among {0.1, 0.2, 0.3, 0.4, 0.5} (Distribution 1; mean=0.3, no fully recessive mutations, Figure 4 and S6), or among {0.0, 0.01, 0.1, 0.2, 0.3, 0.4, 0.5} (Distribution 2; mean=0.22, includes fully recessive mutations, Figure 4 and S7). In these simulations, we aimed at using parameters values close to those observed in mammals, such as human, simulating the evolution of a single panmictic population of *N*=3,125 or *N*=12,500 individuals under a Wright-Fisher model on which mutations occurred at a rate *μ=*1.2*10^-9^ per bp and recombination occurred at a rate *r*=1.2*10^-9^. For each parameter combination (*h*, *s, N,* chromosome type), a simulation was run for 200,000 generations, to allow the population to reach an equilibrium for the number of segregating mutations. At the end of this initialization phase, the nucleotide diversity of populations (π) ranged from 0.000115 (with *s*=-0.01 and distribution 1 used for the dominance coefficients) and 0.000312 (with *s*=-0.001 and distribution 2 used for the dominance coefficients). The non-neutral (with *s*<-1/N) genetic diversity ranged from 0.000040 and 0.000098, indicating that diploid individuals carried on average one deleterious mutation every 10,000-15,000 base pairs. These levels of genetic diversity in terms of deleterious mutations are in the order of magnitude of those estimated in natural populations: for instance, in humans, there is one heterozygous site every 1000bp and about 25% of segregating point mutations have been estimated to be deleterious (Racimo & Schraiber, 2014). Recombination modifiers mimicking inversions of 2Mb and 5Mb were then introduced at the center of the autosome or Y chromosome. We considered inversions either with no intrinsic advantage (s_inv_ = 0.0, meaning their fitness depended solely on the set of mutations they captured) or with an intrinsic fitness advantage (s_inv_ = 0.01). For the latter case, the fitness advantage of inversions was assumed to be codominant, such that homozygotes have a fitness of 1 + sInv and heterozygotes a fitness of 1 + sInv/2. For each parameter combination (*h*, *s*, *N*, chromosome type, size of the chromosomal inversion, fitness effect of the inversion), we ran 100,000 independent simulations starting with the introduction of a single inversion in a single randomly-sampled individual, and we used the same saved initial population for all simulations. We monitored the frequency of these inversion-mimicking mutations during 25,000 generations.

### Simulations of species with haploid phase

To study the effect of the existence of a haploid phase on the accumulation of deleterious mutations and the spread of inversions on autosomes and sex chromosomes (Figure S11), we performed additional simulations, involving 15,000 generations of burn-in in populations of *N*=1000 individuals and the introduction of 10,000 inversions under each combination of parameters in these initial populations. The populations were considered to harbour a single locus with two permanently heterozygous alleles (similarly to the previous situation ii). Every *x* generations, populations experienced a haploid generation, with *x* taking the values 2, 3, 5, 10 or 100. To simulate haploidy, all mutations from one chromosome of each pair (“genome2” in SLiM) were removed, and the dominance coefficient of mutations on the other chromosome (“genome1” of each pair) was set to 1. The recombination rate was set to 0, and mating could only occur between gametes that were derived from the first chromosome of each pair. Therefore, during haploid generations, selection acted only on the first chromosome of each pair, and the second chromosome had no contribution to the following generation. These modifications allowed simulating the occurrence of haploid phases without changing most parameters or the model behavior. Note that, during the haploid phase, the number of individuals remained unchanged, but the number of haploid genomes was divided by two, because the second haploid genome of each pair had no contribution to the following generation.

### Simulations of chromosomal fusion

To study the evolution of chromosomal fusion (Figure S12), we simulated, during 15,000 generations (burn-in) populations of *N*=1000 individuals with two pairs of chromosomes of size 10Mb, one being a mammal-like sex-chromosome with no recombination between X and Y chromosomes between the position 1Mb and 9Mb (chromosome 1), the other being an autosome (chromosome 2). This mimicked the evolution of a population with old sex chromosomes and small pseudo-autosomal regions (1Mb on each chromosome edge). Then, we introduced a fusion-mimicking mutation resulting in the linkage of one sex chromosome (chromosome 1, X or Y) and one autosome (chromosome 2), and suppressing the recombination when heterozygous over 1Mb of the fused side of each chromosome (see figure S11 for a graphical representation). Therefore, these mutations behaved like 2Mb inversions that would also lead to chromosome fusion and result in the extension of the size of the non-recombining region from 8Mb to 10Mb. We tracked the frequency of 10,000 X-autosome and Y-autosome fusion-mimicking mutations for each parameter combination (as was done before for inversion-mimicking mutations).

### Statistical analyses of fixation probabilities

Confidence intervals for fixation probabilities (Figure 3 and S4) were computed assuming binomial sampling with parameter k and t, with k, the number of observed inversion fixation event, and t, the number of inversions simulated (50,000 per parameters set); 95% confidence bounds were extracted as the lower and upper limits of the interval. Differences in inversion fixation between the Y chromosome and autosomes were quantified as the log₂ fold change (log2FC) of fixation probabilities normalized by their neutral expectation. Specifically, we computed Log2[p_Y_/q_Y_/(p_A_/q_A_)] where p_Y_ = k_Y_/t_Y_ and p_A_ = k_A_/t_A_ denote the observed proportions of fixation events on the Y chromosome and autosomes, respectively, and q_Y_ and q_A_ denote their corresponding neutral expectations (2/N and 1/2N, respectively). Uncertainty in fixation probabilities was propagated under a binomial sampling model using a parametric Monte Carlo approach. Fixation probabilities were modeled using the conjugate Beta posterior distribution *Beta*(k + 0.5, n − k + 0.5) (Jeffreys prior). We generated 50,000 posterior draws of p_Y_ and p_A_, calculated the normalized log2FC for each draw, and report the median and 95% credible interval (2.5–97.5% quantiles). Because neutral expectations were treated as fixed quantities, normalization shifts the location of the log2FC distribution without affecting its uncertainty. When no fixation events were observed in either compartment (k_Y_ = k_A_ = 0), the normalized log2FC was set to zero to avoid prior-driven differences arising solely from the Jeffreys prior. This procedure accounts for binomial sampling variance and accommodates asymmetry in uncertainty when fixation events are rare.

### Simulations of sex chromosomes with evolutionary strata

To study more specifically the formation of evolutionary strata on sex chromosomes (Figures 4, 5 and S13), we also simulated the evolution of two 100 Mb chromosomes, one of which carried an XY sex-determining locus at the 50 Mb position, over 115,000 generations (including an initial burn-in of 15,000 generations): individuals could be either XX or XY and could only mate with individuals of a different genotype at this locus. We simulated randomly mating populations of *N*=1000 and *N*=10,000 individuals. Point mutations appeared at a rate of *μ*=10^-9^ per bp, and their individual selection coefficients were determined by sampling a gamma distribution with a mean of −0.03 and with a shape of 0.2; these parameter values were set according to observations in humans (Eyre-Walker & Keightley, 2007; Kim et al., 2017). For each new mutation, a dominance coefficient was chosen from the following values, considered to have uniform probabilities: 0, 0.001, 0.01, 0.1, 0.25, 0.5. At the beginning of each simulation, we randomly sampled *k* genomic positions over the two chromosomes that could be used as inversion breakpoints, with *k* being 10, 100, 1000 or 10,000. After the 15,000 generations of the burn-in period allowing populations to reach an equilibrium in terms of the number of segregating mutations, we introduced *j* inversions in the population in each generation, *j* being sampled from a Poisson distribution of parameter λ=*N*\* k *u_i,_ u_i_ being the inversion rate.

In order to keep the simulation time tractable, we used inversion rates allowing one inversion to occur on average in each generation in the population. For each inversion, the first breakpoint was randomly chosen among the *k* positions, and the second among the potential breakpoint positions less than 20Mb apart on the same chromosome (considering therefore only a subset of the *k* positions for the second breakpoints). Two independent inversions could use the same breakpoints, allowing in particular inversion reversions restoring recombination. If two independent inversions occurred with the same breakpoints on different haplotypes (for instance on the X and on the Y chromosome), we assumed that recombination was restored between these haplotypes, as a reversion would do. The occurrence of partially overlapping inversions (*i.e*., with different breakpoints) on different haplotypes did not restore recombination between these haplotypes. We assumed that inversions subsequently partially overlapped by another inversion or that captured a smaller inversion could not reverse, *i.e*. the restoration of recombination by further inversions, even if using the same breakpoints, was prevented. For *N*=1000 and for each value of *k* (*i.e*., 10, 100, 1000, 10000), we ran 10 simulations (Figure 6). For *N*=10,000, because of computing limitations (each simulation taking about three weeks to run), we only ran one simulation per number of breakpoints.

Simulations were conducted using either GNU Parallel (Tange, 2025) or snakemake (Mölder et al., 2021) and plots were made with ggplot2 (Wickham, 2016) and edited with Inkscape. All scripts are available at git@github.com:PaulYannJay/MutationShelteringTheoryV2.git.

## Acknowledgement

All authors acknowledge the contribution of Emilie Tezenas, co-author of (Jay et al. 2022), who has now left academia and is therefore not an author on this new version of the work. They also thank Ricardo Rodriguez de la Vega, Fanny Hartmann, Jacqui Shykoff, Sylvain Billiard, Janis Antonovics, Olivier Tenaillon, Denis Roze and Diala Abu Awad for insightful discussions and comments on previous draft versions of the manuscript. AV acknowledges support from the chaire program « Mathematical modelling and biodiversity » (Ecole Polytechnique, Museum National d’Histoire Naturelle, Veolia Environnement, Fondation X).

## Author contributions

Original ideas, PJ and TG; deterministic model conception, PJ with input from AV; simulations and data analyses, PJ; interpretation, PJ and TG; manuscript writing, PJ, AV and TG; project management and funding, PJ and TG.

## Data and materials availability

The SLiM and R scripts used to produce all main and supplementary figures are available from GitHub (git@github.com:PaulYannJay/MutationShelteringTheoryV2.git). The numerical outputs of the original simulations from Jay et al. 2022 used to produce the figures in this manuscript are available from Figshare (doi: 10.6084/m9.figshare.19704457). The output from the new simulations will be uploaded on Figshare upon publication. Details concerning the mathematical modeling are available in the appendix.

