## Supplementary material for "Stepwise expansion of recombination suppression on sex chromosomes and other supergenes through lower load advantage and deleterious mutation sheltering"

**Affiliations:**
[1]Laboratoire d'Ecologie Alpine, CNRS, Univ. Grenoble Alpes, Univ. Savoie Mont Blanc, 38000 Grenoble, France
[2]Université Paris Cité, CNRS, MAP5, F-75006 Paris, France
[3]Ecologie Société Evolution, IDEEV, Bâtiment 680, 12 route RD128, 91190 Gif-sur-Yvette, France

**This document contains:**
Text S1
Figures S1-14

***Infinite population deterministic model (Figure S1 & S8).***

If we neglect mutation accumulation following inversion occurrence (*i.e.*, if the fitness of inverted and non-inverted segments remains unchanged through time), the trajectory and fate of an inversion in Y chromosomes and in autosomes can be readily quantified by a deterministic model as follows.

We consider the discrete-time evolution of an infinite size, randomly mating population experiencing only deleterious recessive mutations, with heterozygotes and homozygotes suffering from a *1-hs* and *1-s* reduction in fitness, respectively. At all *n* sites, mutations are at the same mutation-selection equilibrium frequency, denoted *q*. We used non-approximated values of *q* as derived by Crow (1970):

$$q = \frac{h(1+u)}{2(2h-1)}\left[1-\sqrt{1-\frac{4(2h-1)u}{sh^2(1+u)^2}}\right]$$

We assume that all sites are independent. The number of mutations carried by a chromosomal segment of length *n* then follows a binomial distribution of parameters (*n, q*). We follow the frequency of an inversion I of size *n,* considering that it captures *m* mutations and that it appears in a population carrying only non-inverted segments. The mean fitness of a non-inverted homozygote can be computed as follows:

$$W_{NN} = (1-2q(1-q)hs-q^2s)^n$$

Note that in the parameter regimes where we can make the approximation $q \approx u/hs$ with $q << 1$, we have $\overline{W_{NN}} \approx (1-2u)^n$, in accordance with the mutation load theory (Agrawal & Whitlock, 2012). Similarly, an individual who is heterozygous for an inversion with *m* mutations has a mean fitness of

$$W_{NI} = (q(1-s)+(1-q)(1-hs))^m(q(1-hs)+1-q)^{n-m}$$

Assuming that $q << 1$, Nei et al., 1967 considered that individuals heterozygous for an inversion had no homozygous mutations, so that $W_{NI} \approx (1-hs)^m(1-hs)^{nq} \approx (1-hs)^{nq+m}$. Note that we use non-approximated values in our computations, in order to treat all parameter regimes in a similar way. An individual who is homozygous for a segment I with *m* mutations is homozygous for all these mutations. Its fitness can therefore be expressed as $W_{II} = (1-s)^m$.

The inversion frequency trajectory can be determined with a simple two-locus two-allele model. Focusing on inversions partly linked to a permanently heterozygous allele in a XY system, the changes in frequency of inversions on the Y chromosome, on the X chromosome or on autosomes between generations $t$ and $t+1$ are described by

$$
\begin{aligned}
F_{XIm}^{t+1} &= \frac{F_{XIf}^{t}\left(W_{II}F_{YI}^{t} + W_{NI}F_{YN}^{t}\right) + rW_{NI}D^{t}}{\overline{W_m^t}}, \\
F_{XNm}^{t+1} &= \frac{F_{XNf}^{t}\left(W_{NN}F_{YN}^{t} + W_{NI}F_{YI}^{t}\right) - rW_{NI}D^{t}}{\overline{W_m^t}}, \\
F_{XIf}^{t+1} &= \frac{F_{XIm}^{t}F_{XIf}^{t}W_{II} + \frac{1}{2}W_{NI}[F_{XIm}^{t}F_{XNf}^{t} + F_{XIf}^{t}F_{XNm}^{t}]}{\overline{W_f^t}}, \\
F_{XNf}^{t+1} &= \frac{F_{XNm}^{t}F_{XNf}^{t}W_{NN} + \frac{1}{2}W_{NI}[F_{XNm}^{t}F_{XIf}^{t} + F_{XNf}^{t}F_{XIm}^{t}]}{\overline{W_f^t}}, \\
F_{YI}^{t+1} &= \frac{F_{YI}^{t}(F_{XIf}^{t}W_{II} + F_{XNf}^{t}W_{NI}) - rW_{NI}D^{t}}{\overline{W_m^t}}, \\
F_{YN}^{t+1} &= \frac{F_{YN}^{t}(F_{XNf}^{t}W_{NN} + F_{XIf}^{t}W_{NI}) + rW_{NI}D^{t}}{\overline{W_m^t}}, \\
F_{XI} &= \frac{2}{3}F_{XIf} + \frac{1}{3}F_{XIm}, \\
F_{I} &= \frac{1}{4}F_{YI} + \frac{3}{4}F_{XI}
\end{aligned}
$$

(Equation 1)

where $F_{YI}^{t}$ is the frequency of inversions in the population of Y chromosomes at time *t*, $F_{XIf}^{t}$ is the frequency of the inversion on the X chromosome in females (respectively $F_{XIm}^{t}$ in males), *r* is the rate of recombination between the inversion and the sex-determining locus, *D* is their linkage disequilibrium (so that $D^{t} = F_{XNf}^{t}F_{YI}^{t} - F_{XIf}^{t}F_{YN}^{t}$) and $\overline{W_m^t}$ is the mean male fitness. When *r*=0.5, this system of equations describes the evolution of inversions on autosomes. When the inversion captures the male-determining allele, *r*=0, $F_{XIm} = F_{XIf} = 0$, and $F_{XNf} = F_{XNm} = 1$ at any time *t*. The equations for $F_{YI}^{t}$, the frequency of inversions capturing the male-determining allele then reduce to

$$\mathrm{F}_{YI}^{t+1} = F_{YI}^{t}\frac{W_{NI}}{\overline{W_m^t}},$$

$$\mathrm{F}_{YN}^{t+1} = F_{YN}^{t}\frac{W_{NN}}{\overline{W_m^t}}.$$

with $\overline{W_m^t} = F_{YN}^t W_{NN} + F_{XI}^t W_{NI}$. Substituting $F_{YN}$ with $1 - F_{YI}$ and neglecting $F_{YI}^t$, we have

$$\Delta F_{YI} = \frac{F_{YI}^t(1 - F_{YI}^t)\,(W_{NI} - W_{NN})}{\overline{W_m^t}}$$

The search for values of $F_{YI}$ such that $\Delta F_{YI} = 0$ readily gives two equilibria: 0 and 1. Since $\overline{W_m^t} > 0$ and $0 \leq F_{YI} \leq 1$, we can conclude that:

$$\begin{array}{ll} \text{If } W_{NI} > W_{NN}, & F_{YI} \rightarrow 1, \\ \text{If } W_{NI} < W_{NN}, & F_{YI} \rightarrow 0 \end{array}$$

In the case of an inversion appearing on an autosome ($r$=0.5), $F_{XIm} = F_{XIf} = F_{YI}$ and $\overline{W_m} = \overline{W_f}$, so that the change in inversion frequency in the population is:

$$F_I^{t+1} = \frac{(F_I^t)^2 W_{II} + F_I^t F_N^t W_{NI}}{\overline{W^t}},$$

and

$$\Delta F_I = \frac{F_I}{\overline{W}}\Big(F_I^2\,(2W_{NI} - W_{II} - W_{NN}) + F_I\,(W_{II} + 2W_{NN} - 3W_{NI}) + W_{NI} - W_{NN}\Big) = \frac{F_I}{\overline{W}} \times P(F_I)$$

with $P(X) = X^2\,(2W_{NI} - W_{II} - W_{NN}) + X\,(W_{II} + 2W_{NN} - 3W_{NI}) + W_{NI} - W_{NN}$.

The equilibria are 0 and the roots of the polynomial *P*, which are 1 and $\frac{(W_{NI} - W_{NN})}{(2W_{NI} - W_{II} - W_{NN})}$. We thus have:

$$\begin{array}{ll} \text{If } W_{NI} > W_{NN} \text{ and } W_{II} > W_{NI}, & F_I \rightarrow 1, \\ \text{If } W_{NI} > W_{NN} \text{ and } W_{II} < W_{NI}, & F_I \rightarrow \dfrac{W_{NN} - W_{NI}}{W_{II} + W_{NN} - 2W_{NI}} \\ \text{If } W_{NI} < W_{NN}, & F_I \rightarrow 0. \end{array}$$

Therefore, as expected inversion equilibrium frequencies depend on the relative fitness of homozygotes and heterozygotes for the inversion and of the non-inverted homozygotes. We thus derive the conditions for the inversion to be favoured or disfavoured as a function of the number of mutations captured by the inversion (*m*)*.* A straightforward computation:

$$\begin{array}{lll} W_{II} > W_{NI} & \text{if and only if} & m < \beta_1(q,h,s) \times n, \\ W_{NI} > W_{NN} & \text{if and only if} & m < \beta_2(q,h,s) \times n \end{array}$$

with

$$\beta_1(q,h,s) = \frac{\ln(q(1-hs)+1-q)}{\ln\left(\frac{(1-s)(q(1-hs)+1-q)}{q(1-s)+(1-q)(1-hs)}\right)} = \frac{\ln(1-qhs)}{\ln\left(\frac{(1-s)(1-qhs)}{1-hs-qs(1-h)}\right)}, \quad \text{(Equation 2)}$$

$$\beta_2(q,h,s) = \frac{\ln\left(\frac{1-2q(1-q)hs-q^2s}{q(1-hs)+1-q}\right)}{\ln\left(\frac{q(1-s)+(1-q)(1-hs)}{q(1-hs)+1-q}\right)}. \quad \text{(Equation 3)}$$

Assuming *q*<<1 and *s*<<1, these quantities can be approximated by $\beta_1 \approx q\frac{h}{1-h}$ and $\beta_2 \approx q$. When $qhn/(1-h) < m < nq$, inversions should thus go to fixation on Y chromosomes and stabilize at intermediate frequency on autosomes. When $m < qhn/(1-h)$, inversions should go to fixation on autosomes and on the Y chromosome. Observe that the closer *h* is to ½ (*i.e.*, the scenario without dominance), the smaller the difference between the thresholds $\beta_1$ and $\beta_2$ is. When *h*=0.5, $\beta_1 = \beta_2$, inversions should therefore go to fixation on autosome when they have fewer mutations than average, as they then have no homozygous disadvantage preventing their fixation. In contrast, when *h* is small, $\beta_1$ is significantly smaller than $\beta_2$, showing that the condition for heterozygotes to be favoured over non-inverted homozygotes ($W_{NI} > W_{NN}$) is much easier to meet than the condition for inversion homozygotes to be favoured over inversion heterozygotes ( $W_{II} > W_{NI}$): inversions are therefore much more likely to be maintained at intermediate frequencies on autosomes than to fix.

We used these equilibrium frequencies as functions of *m* to compute the expected equilibrium frequency of inversions that can occur in the genome ($F_{equ}$). To do so, we sum, for all $m \in \{0, \dots, n\}$ the equilibrium frequency of inversions ($F_{equ,m}$) weighted by their occurrence probability ($P_m$). For inversions on the Y chromosome, we therefore have:

$$\begin{aligned}[F_{equ}] &= \sum_{m=0}^{n} P_m \times F_{equ,m} \\ &= \sum_{m=0}^{n} \binom{n}{m} q^m (1-q)^{n-m} F_{equ,m} \\ &= \sum_{m=0}^{\lfloor nq \rfloor} \binom{n}{m} q^m (1-q)^{n-m}.\end{aligned} \qquad \text{(Equation 4)}$$

For inversions on autosomes, we obtain, using the approximate values for $\beta_1$ and $\beta_2$ derived earlier in the case q<<1 and s<<1:

$$[F_{equ}] = \sum_{m=0}^{\lfloor nq\frac{h}{1-h} \rfloor} \binom{n}{m} q^m (1-q)^{n-m} + \sum_{m=\lfloor nq\frac{h}{1-h} \rfloor+1}^{\lfloor nq \rfloor} \binom{n}{m} q^m (1-q)^{n-m} \frac{W_{NI} - W_{NN}}{2W_{NI} - W_{NN} - W_{II}}$$

(Equation 5)

The expected equilibrium frequency of less-loaded inversions is therefore the expected equilibrium frequency of all inversions divided by the probability of occurrence of less-loaded inversions:

$$\begin{aligned}[F_{equ}^{less-loaded}] &= \frac{[F_{equ}]}{\sum_{m=0}^{\lfloor nq \rfloor} P_m} \\ &= \frac{[F_{equ}]}{\sum_{m=0}^{\lfloor nq \rfloor} \binom{n}{m} q^m (1-q)^{n-m}}\end{aligned} \qquad \text{(Equation 6)}$$

Readers interested in further details of the derivation are referred to the appendix of our PLoS Biology paper (now retracted but still available), available at: https://journals.plos.org/plosbiology/article?id=10.1371/journal.pbio.3001698#sec018

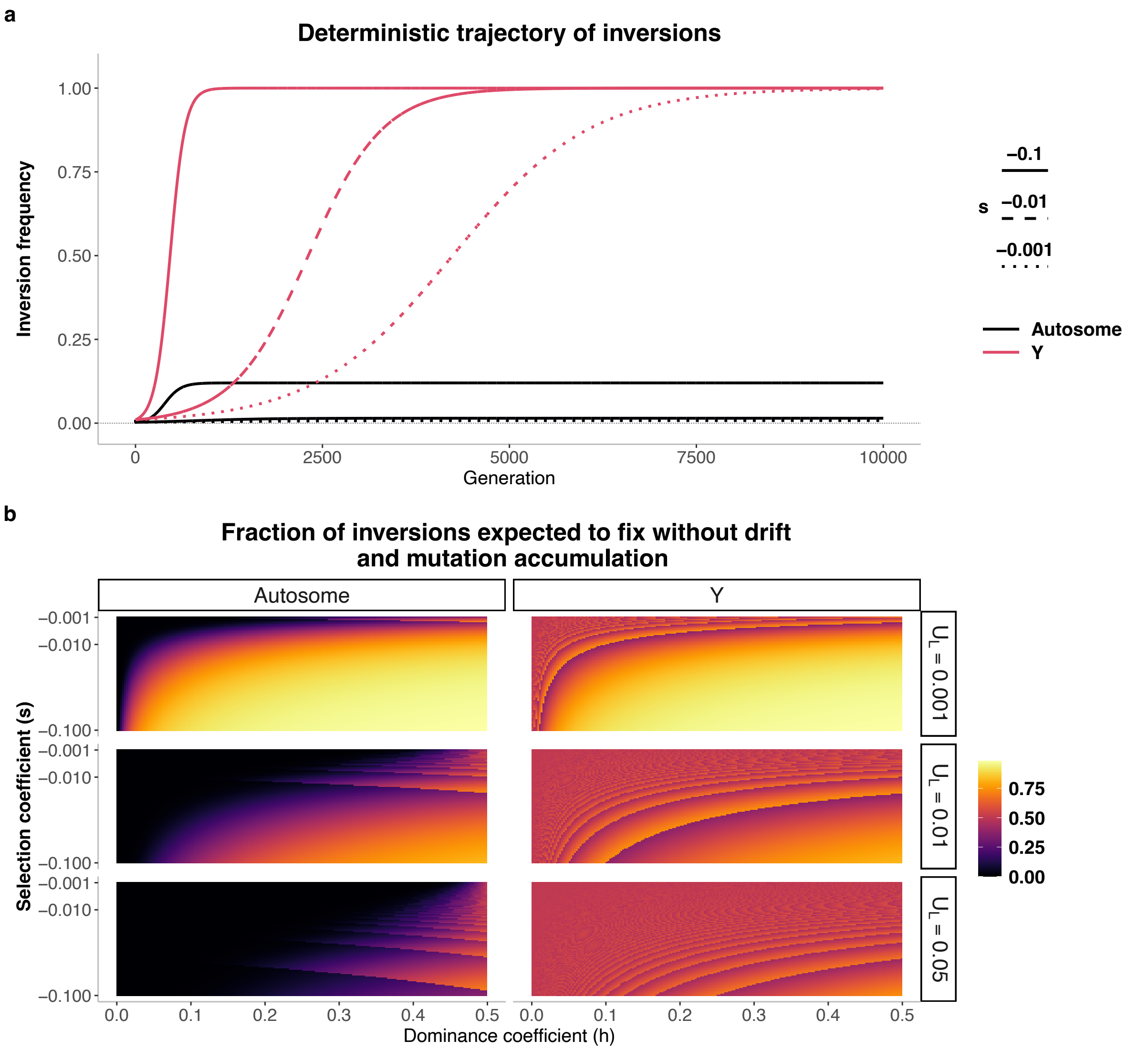


**Figure S1 | Without drift, less-loaded inversions are more likely to fix on Y chromosomes than on autosomes.**

**a.** Deterministic change in frequency for 2Mb inversions, either capturing the sex-determining allele on the Y chromosome or occurring on an autosome. For clarity, the frequency shown for Y-linked inversions refers to their frequency within the population of Y chromosomes. Inversions are assumed to carry 5% fewer mutations than the population average ($m = 0.95 \times nq$), with mutations at mutation–selection equilibrium ($\mu = 10^{-8}$, $U_L = 0.02$, $h = 0.1$). **b.** Proportion of inversions that are expected to fix in deterministic simulations without mutation accumulation after inversion formation—defined here as the fraction of inversions that confer a selective advantage over the course of the simulation. For Y-linked inversions, this corresponds to inversions carrying fewer mutations than the population average ($m < nq$). For autosomal inversions, this corresponds to inversion with $m < qnh/(1-h)$, where $q$ is the equilibrium frequency of deleterious alleles. See supplementary text 1 for derivation details.

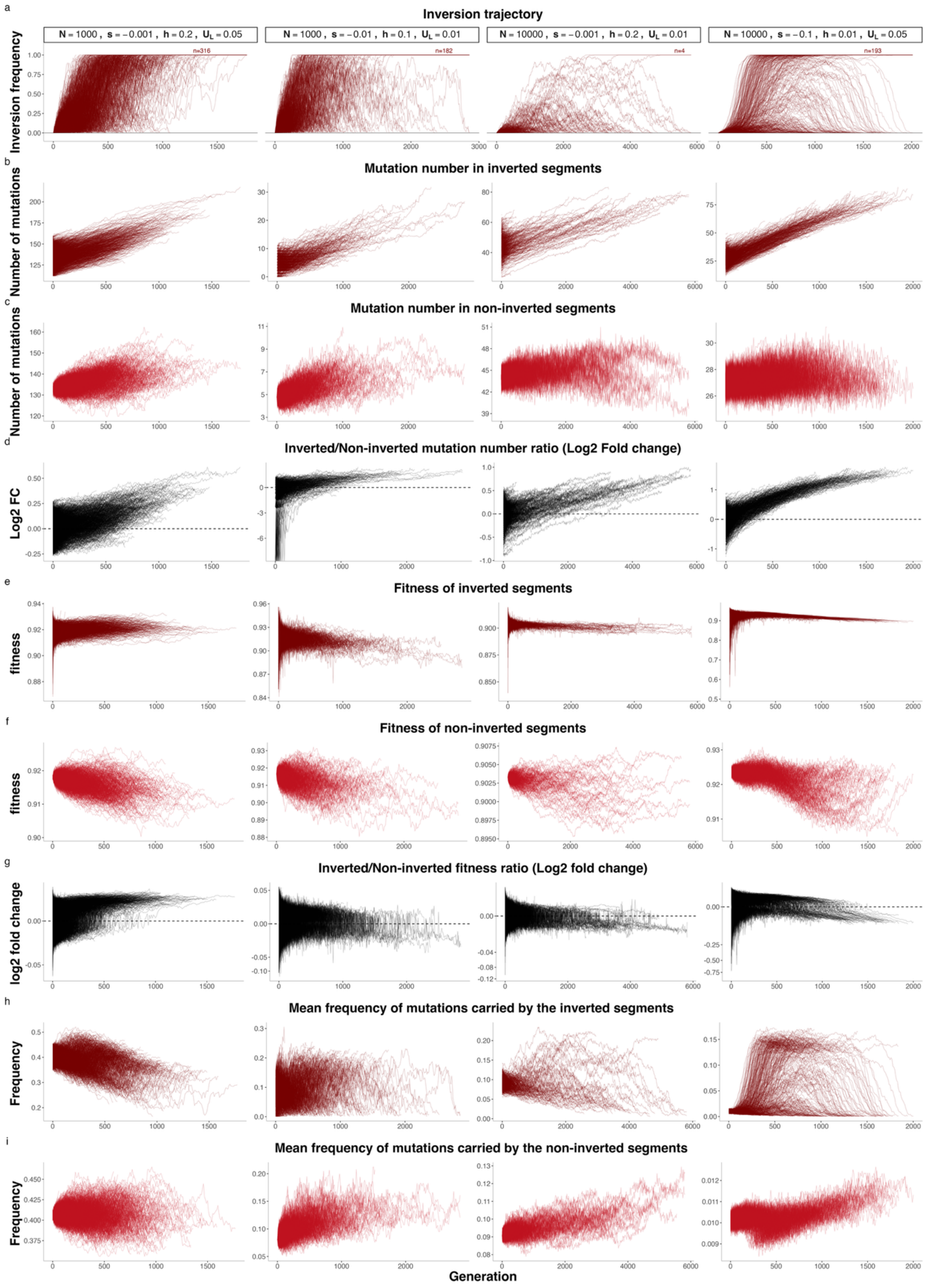


**Figure S2 | Trajectories of inversions in individual-based simulations on Y chromosomes.**

Four combinations of parameters are shown. For each combination, the trajectories of 50,000 inversions are displayed until the generation at which all inversions are either fixed or lost. **a**, Change in inversion frequency. **b**, Number of mutations in the inverted segment. **c**, Number of mutations in the non-inverted segments. **d**, Log_2 ratio of mutations in inverted versus non-inverted segments. **e**, Average fitness of individuals carrying inverted segments, calculated using SLiM's cachedFitness function. **f,** Average fitness of individuals carrying non-inverted segments, calculated using SLiM's cachedFitness function. **g**, Log_2 ratio of the fitness of individuals carrying inverted versus non-inverted segments. **h**, Average population frequency of mutations occurring in inverted segments. **i**, Average population frequency of mutations occurring in non-inverted segments.

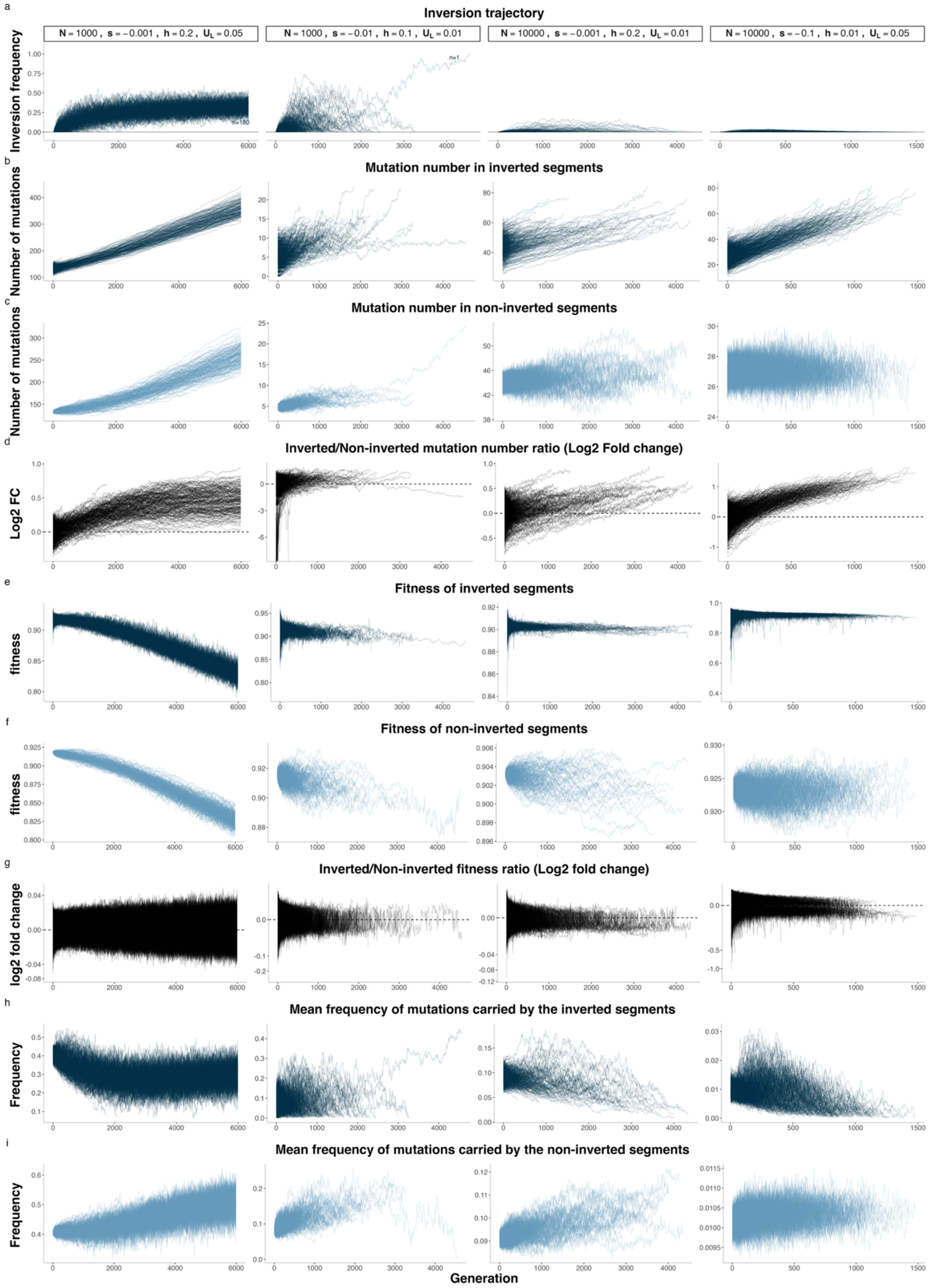


**Figure S3 | Trajectories of inversions in individual-based simulations on autosomes.**

Four combinations of parameters are shown. For each combination, the trajectories of 50,000 inversions are displayed until the generation at which all inversions are either fixed or lost. **a**,

Change in inversion frequency. **b**, Number of mutations in the inverted segment. **c**, Number of mutations in the non-inverted segments. **d**, Log_2 ratio of mutations in inverted versus non-inverted segments. **e**, Average fitness of individuals carrying inverted segments, calculated using SLiM's cachedFitness function. **f,** Average fitness of individuals carrying non-inverted segments, calculated using SLiM's cachedFitness function. **g**, Log_2 ratio of the fitness of individuals carrying inverted versus non-inverted segments. **h**, Average population frequency of mutations occurring in inverted segments. **i**, Average population frequency of mutations occurring in non-inverted segments.

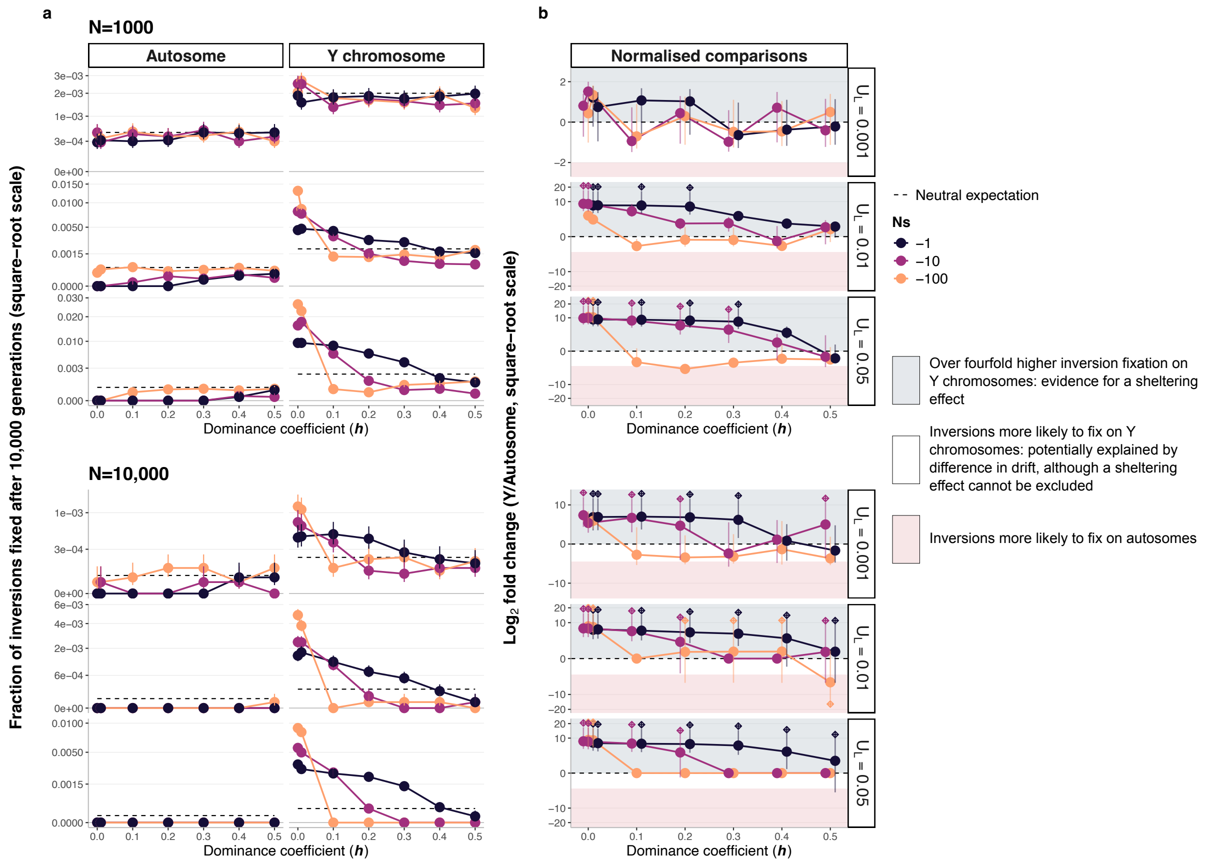


**Figure S4 | Less-loaded inversions are more likely to fix on Y chromosomes than on autosomes.**

This figure is identical to Figure 3, except that Y-linked inversions were simulated for 6N/4 generations instead of 6N generations, thereby accounting for differences in effective population size between autosomes and Y chromosomes.. **a.** Proportion of inversions that reached fixation after 10,000 generations in stochastic simulations with N = 1,000 and N = 10,000 across different combinations of parameter values (N, N*s, h, $U_L$) after 6N generations in autosome and 6N/4 generation in Y chromosome. For each parameter combination, 50,000 inversions capturing a random genomic fragment were simulated. Dashed lines represent the neutral expectation, i.e. 2/N for Y chromosomes and 1/(2N) for autosomes. In contrast to Figure 3c in Jay et al. (2022), all inversions are shown here, rather than only the loaded inversions that survived the first 20 generations. Error bars represent 95% binomial confidence intervals; none are shown when no fixation occurred. **b.** $\log_2$ ratio of the normalized fixation probabilities for Y-linked versus autosomal inversions. Absolute fixation probabilities (shown in panel a) were normalized by dividing them by their respective neutral expectations. Values above 0 indicate that inversions had more than fourfold higher absolute fixation probability on Y chromosomes (consistent with a sheltering effect), whereas values below −2 indicate higher absolute fixation probability on autosomes. To represent cases in which no autosomal or Y-linked inversion fixed in the simulations—which would otherwise produce an infinite ratio—we assumed that 0.5 inversions fixed in those cases. These situations are indicated by diamonds above or below the corresponding points. Diamonds above 0 indicate scenarios where no inversion fixed on autosomes, whereas diamonds below 0 indicate that no inversion fixed on the Y chromosome.  Error bars show the 95% credible interval of the log2 fold change estimated from Monte Carlo sampling (see Methods).

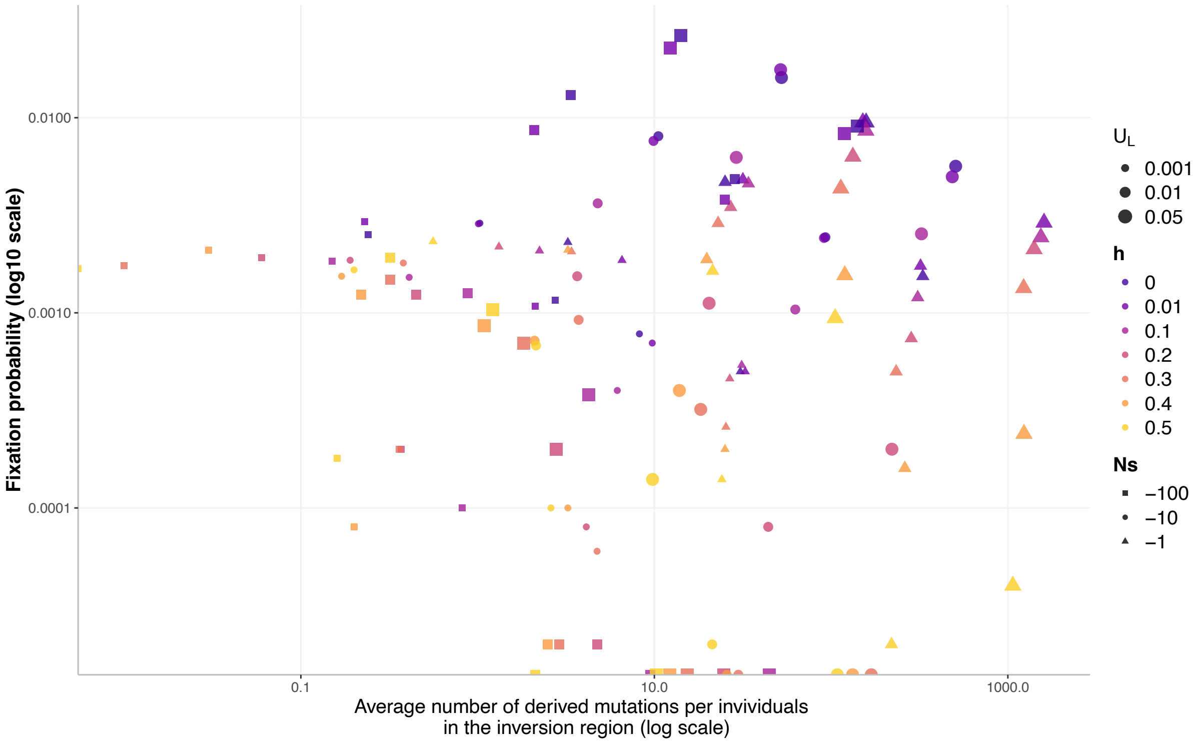


**Figure S5 | Inversion fixation probability as function of the number of derived mutations in the inverted regions.**

The same simulations shown in Figure 3 are displayed here.

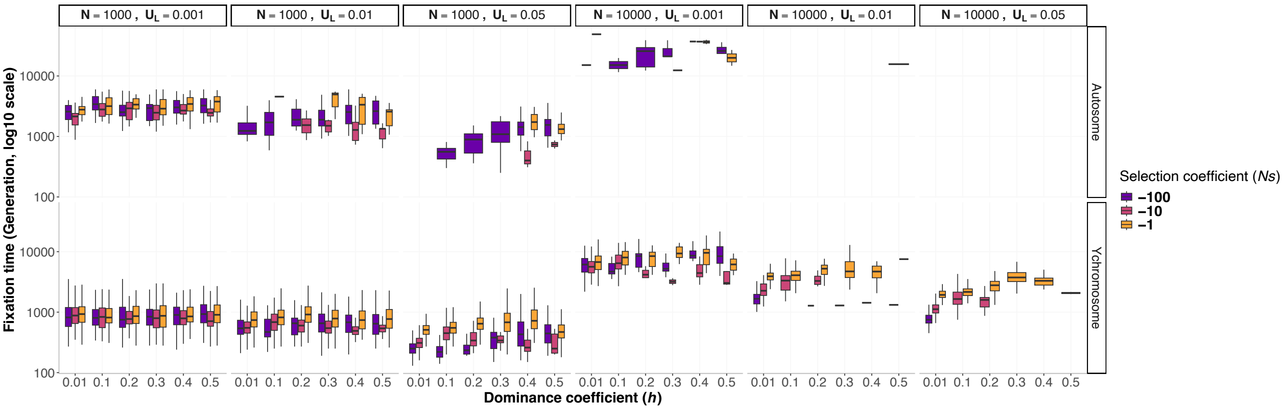


**Figure S6 | Inversion fixation time.**

The plot displays the fixation time of the simulations shown in Figure 3.

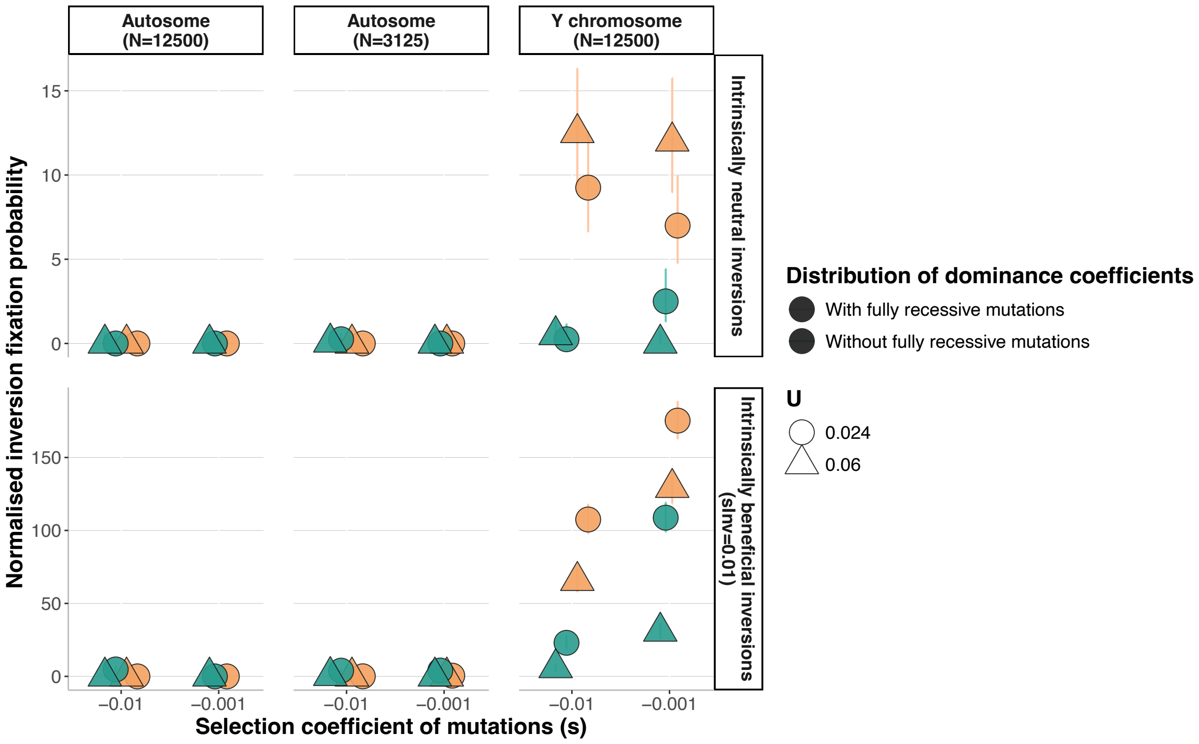


**Figure S7 | The sheltering effect facilitates inversion fixation on Y-like sex chromosomes when segregating mutations have different dominance coefficient and when inversions are intrinsically beneficial.**

Proportion of inversions that reached fixation after 6*N* generations in stochastic simulations divided by their respective neutral fixation probability (*i.e.*, normalized probability), across different combinations of parameter values with *N*=12,500 or *N*=3,125. This figure reports simulations where mutations had their fitness coefficients sampled from a gamma distribution of mean -0.001 or -0.01, and their dominance coefficient randomly sampled with uniform probabilities among either {0.1, 0.2, 0.3, 0.4, 0.5} (mean=0.3, no fully recessive mutations), or {0.0, 0.01, 0.1, 0.2, 0.3, 0.4, 0.5} (mean=0.22, with fully recessive mutations). To show that the sheltering effect can act even in the absence of the lower-load advantage, we performed simulations of inversions benefiting from an intrinsic selective advantage ($s_{Inv}$=0.01), in addition to the advantage or disadvantage conferred by their relative mutation load. A total of 100,000 inversions of 2Mb and 5Mb were studied for each combination of parameter values. In addition to the simulations with *N*=12,500, we also performed simulations with an autosomal population of size equivalent to the effective population size of the Y chromosome, *i.e.*, 3,125 (¼ of 12,500), as another approach to control for difference in population size between Y chromosome and autosome.

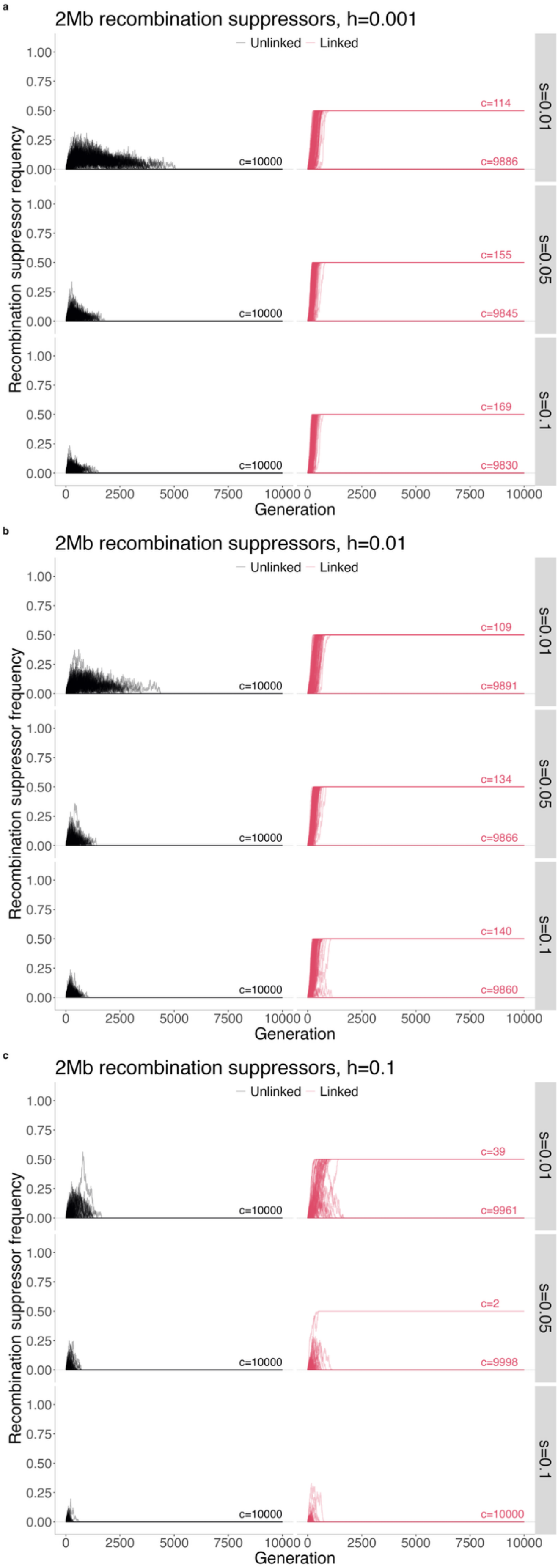


**Figure S8. Evolution of 2Mb recombination suppressors ($U_L$=0.02) in linkage with a locus with two permanently heterozygous alleles in stochastic simulations during 10,000 generations.**
For each parameter combination, 10,000 recombination suppressors of 2Mb were simulated. These recombination modifiers suppress recombination within the segment in which they reside, both when heterozygous and when homozygous. These recombination suppressors were either fully linked (*r*=0.0) or fully unlinked (*r*=0.5) to a locus with two permanently heterozygous alleles; recombination suppressors fully linked to one of the two permanently heterozygous alleles can spread to a maximum frequency of 50%. At the end of the simulation, all recombination suppressors display either a 0.0 or a 0.5 frequency, meaning they are either lost or fixed to a given permanently heterozygous allele. The number of recombination suppressors in each state at the end of the simulation is indicated above lines (« c=... »). Populations of *N*=1000 individuals were simulated.

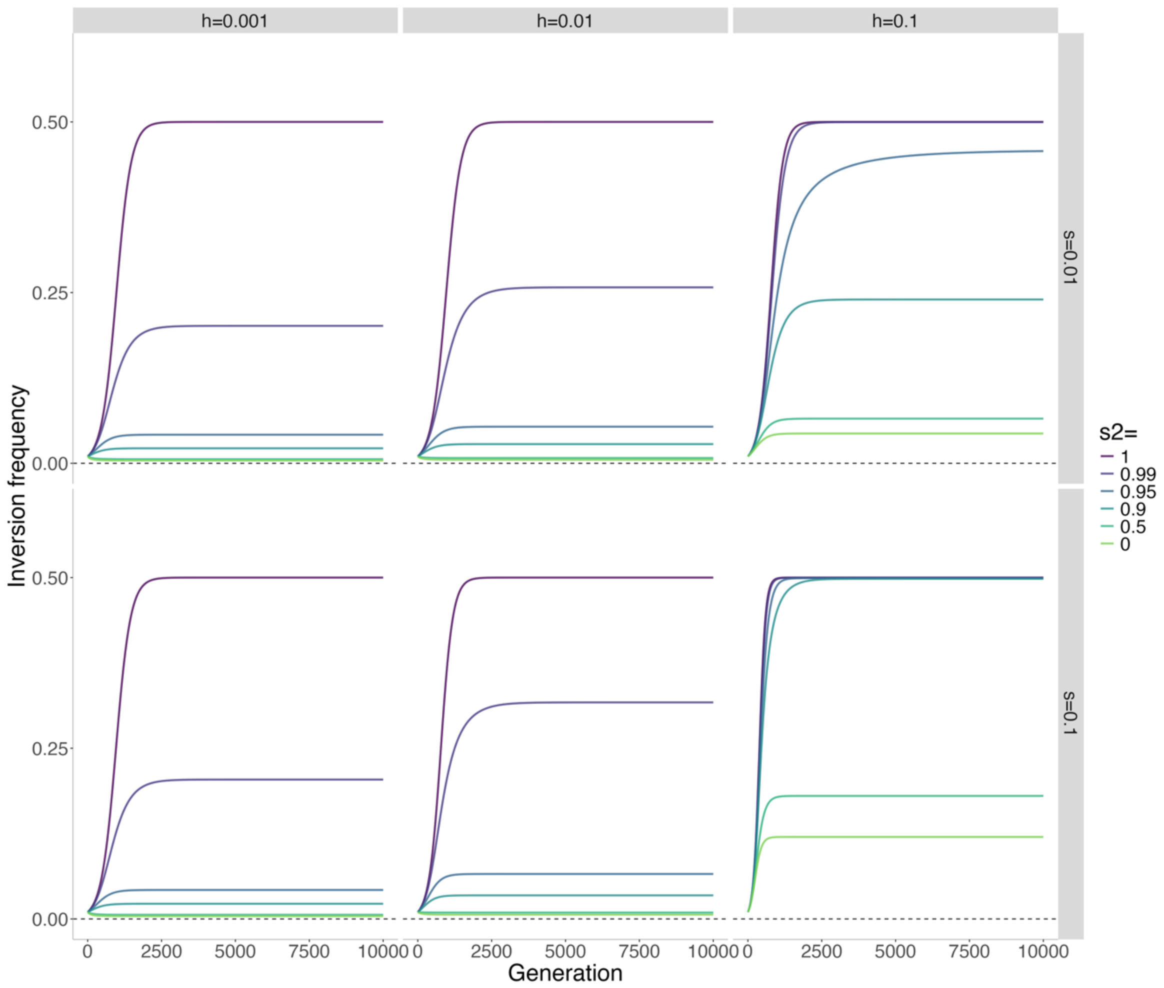


**Figure S9. Deterministic trajectory of inversion frequency when in linkage with an overdominant locus with no permanently heterozygous alleles.**

To save computation time, we only analysed the evolution of overdominant inversions in a deterministic model without mutation accumulation (similar to Figure S1a). In this model, inversions appear in full linkage (*r*=0.0) with one the two alleles at an overdominant locus where both homozygotes have their fitness reduced by a value $s_2$, such the fitness of individual homozygous for the non-inverted segment is $W_{NN} = (1 - 2q(1-q)hs - q^2 s)^n - s_2$ and the fitness of inversion homozygotes is $W_{II} = (1-s)^m - s_2$ (See supplementary text 1). When $s_2$ approaches 1, homozygotes experience a stronger reduction in fitness and consequently become rarer. The plot displays the trajectory of inversions depending on the selection and dominance coefficients of the mutations segregating in the genome and the strength of selection acting on the overdominant locus (with no permanently heterozygous alleles). Mutations were considered to be at mutation-selection equilibrium frequency (*q*)with a mutation rate of *u*=10-8. The figure illustrates the case of *n*=2Mb inversions carrying a number of mutations 20% lower than the population average (i.e. *m*=[0.8**nq*]).

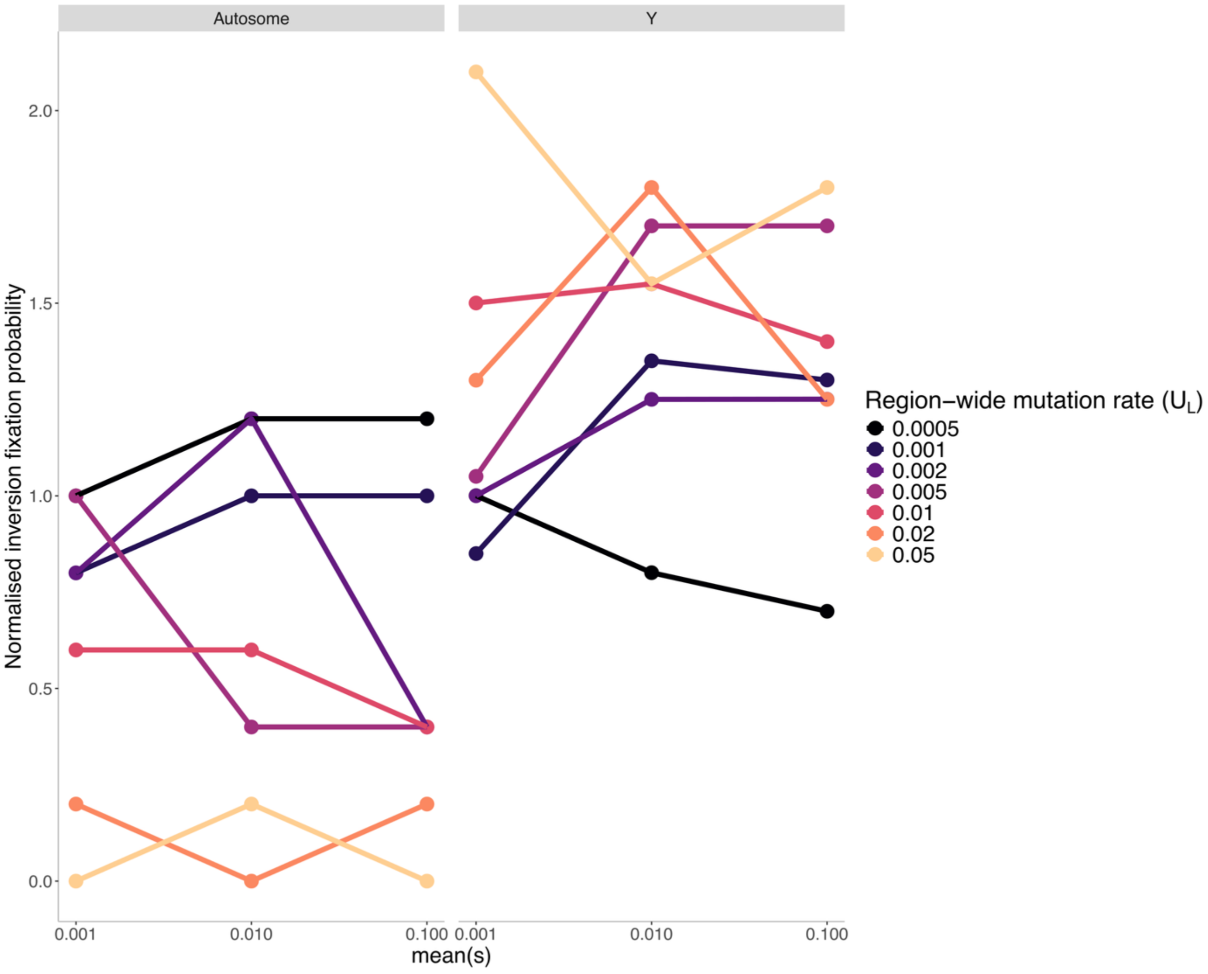


**Figure S10 | Normalised proportions of inversions in stochastic simulations that were fixed after 10,000 generations, for different parameter value combinations with *N*=1000 and with variable mutation fitness effects.**

Similar to Figure 3 and Figure 4 but considering that mutations segregating in the genome have their fitness effects drawn from a gamma distribution with a shape of 0.2, and their dominance coefficient *h* randomly sampled among 0, 0.001, 0.01, 0.1, 0.25, 0.5 with uniform probabilities. Simulations were run considering that the mean of the selection coefficient values (mean of the gamma distribution) was either 0.001, 0.01 or 0.1. The absolute inversion fixation probabilities were normalized by dividing them by their respective neutral expectations.

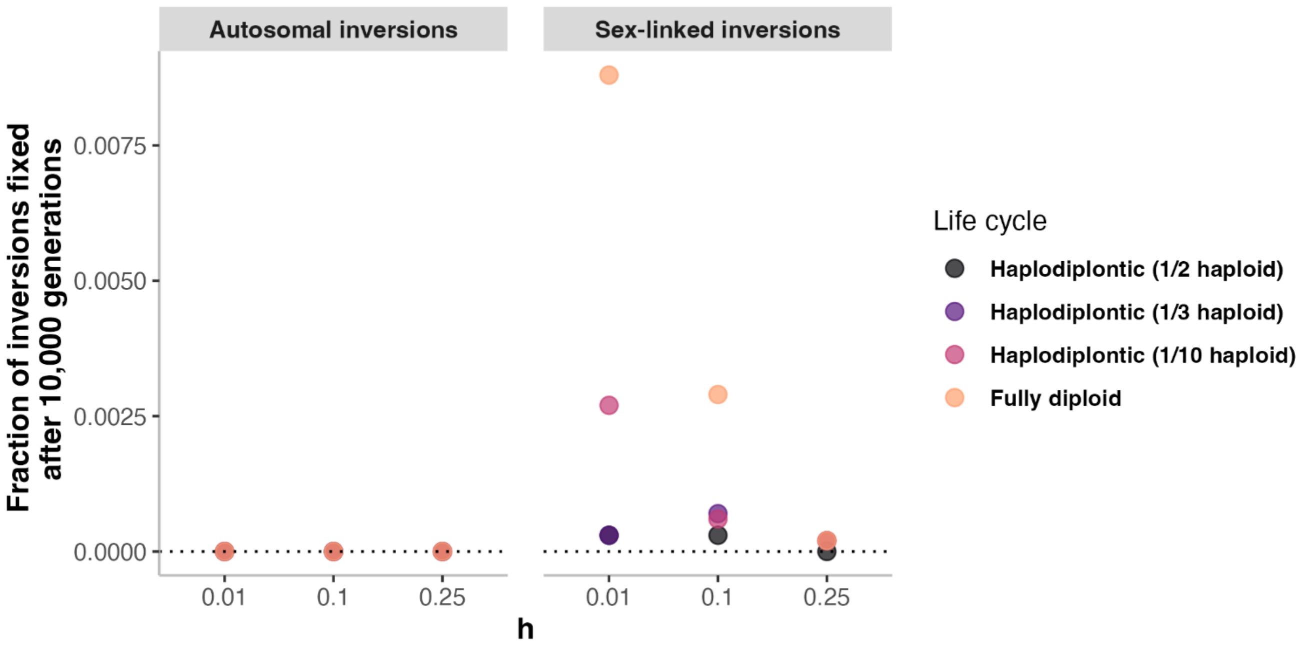


**Figure S11 | Proportions of inversions in stochastic simulations that were fixed after 10,000 generations, for different parameter value combinations, with *N*=1000 and in populations with haplodiplontic life cycles.**

Similar to Figure 4 but considering species with different life cycles. Four life cycles were considered: fully diploid (as used throughout this study), and haplodiplontic with occurrence of a haploid phase every *x* generation, with *x* being 2, 3 or 10. Simulations were performed with *N*=1000, $U_L$=0.01, *s*=-0.01. This shows that inversions are less likely to spread and fix in species with extended haploid phases, as this type of life cycles lead to a more efficient purge of deleterious recessive mutations.

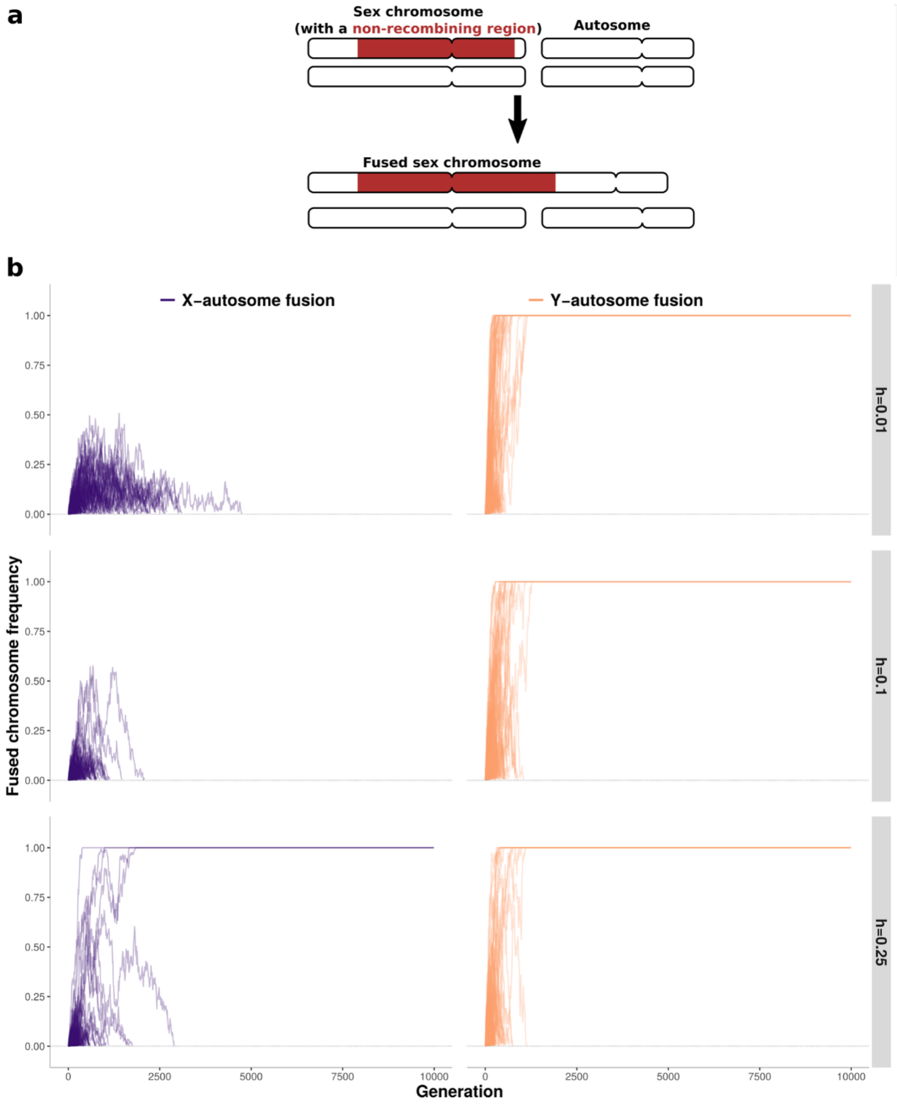


**Figure S12 | Evolution of fusion-mimicking mutations in stochastic simulations during 10,000 generations.**

**a**, Graphical representation of the chromosome fusions simulated. The fusion-mimicking mutations result in the linkage of two chromosomes of each pair and in the suppression of recombination in heterozygotes over 1Mb in the fused side of each chromosome. Therefore, these mutations behave as 2Mb inversions that would also lead to chromosome fusion. **b**, Similar to Figure S2a but considering fusion-mimicking mutations instead of inversions. Simulations were performed with $N$=1000, $u$=5.10$^{-9}$ and $s$=-0.01. For each parameter combination, 10,000 fusion-mimicking mutations were simulated in a population harboring a pair of autosomes and a pair of XY-like non-recombining sex chromosomes (females being XX and males XY) with a 1Mb recombining region on each sex-chromosome side (mimicking pseudo-autosomal regions). Only fusions not lost after 20 generations are displayed. The displayed frequency of the fused chromosomes is their frequency among the chromosomes of the same type, *e.g.* a Y-autosome fusion at 1.0 frequency means that all Y chromosomes are fused with an autosome.

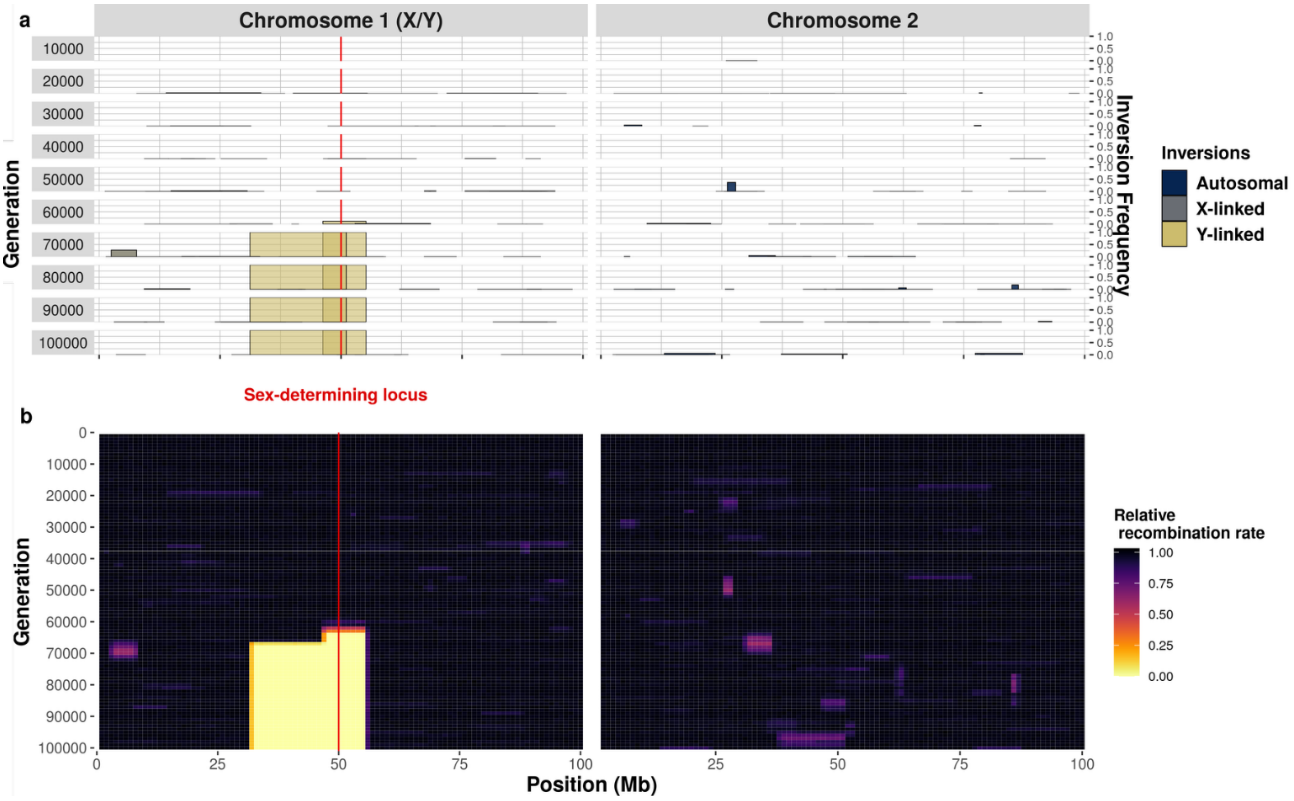


**Figure S13 | Successive accumulation of inversions around a male-determining allele in an XY system, leading to the formation of non-recombining sex chromosomes.**

Similar to Figure 5 but displaying the result of another simulation with a population of *N*=10,000 individuals, each with two pairs of 100 Mb chromosomes, during 100,000 generations. Chromosome 1 harbors an X/Y sex-determining locus at 50 Mb (individuals are XX or XY). Each generation, one inversion appears on average in the whole population, in an individual sampled uniformly at random, with the two recombination breakpoints sampled uniformly at random among *k*=100 potential breakpoints. **A,** Overview of chromosomal inversion frequency and position for 10 different generations. Square width represents inversion position and square height inversion frequency. Inversions appearing on the Y chromosome are depicted in yellow, those appearing on the X chromosomes in gray. The colors are not entirely opaque, so that regions with overlapping inversions appear darker. Loss of previously fixed inversions are due to beneficial reversion occurrence and selection. **B,** Changes in the relative rate of recombination over the entire course of the simulation. The numbers of recombination events occurring at each position (binned in 1Mb windows) are recorded at the formation of each offspring, across all homologous chromosomes in the population. To illustrate the evolution of recombination suppression between the sex chromosomes, only recombination events between the X and the Y chromosomes are shown for chromosome 1 (*i.e.*, not the recombination events between the two X chromosomes in females). Unlike chromosome 1, chromosome 2 harbors no permanently heterozygous allele. All inversions on this chromosome suffer from homozygosity disadvantage and very few inversions therefore become fixed on chromosome 2.

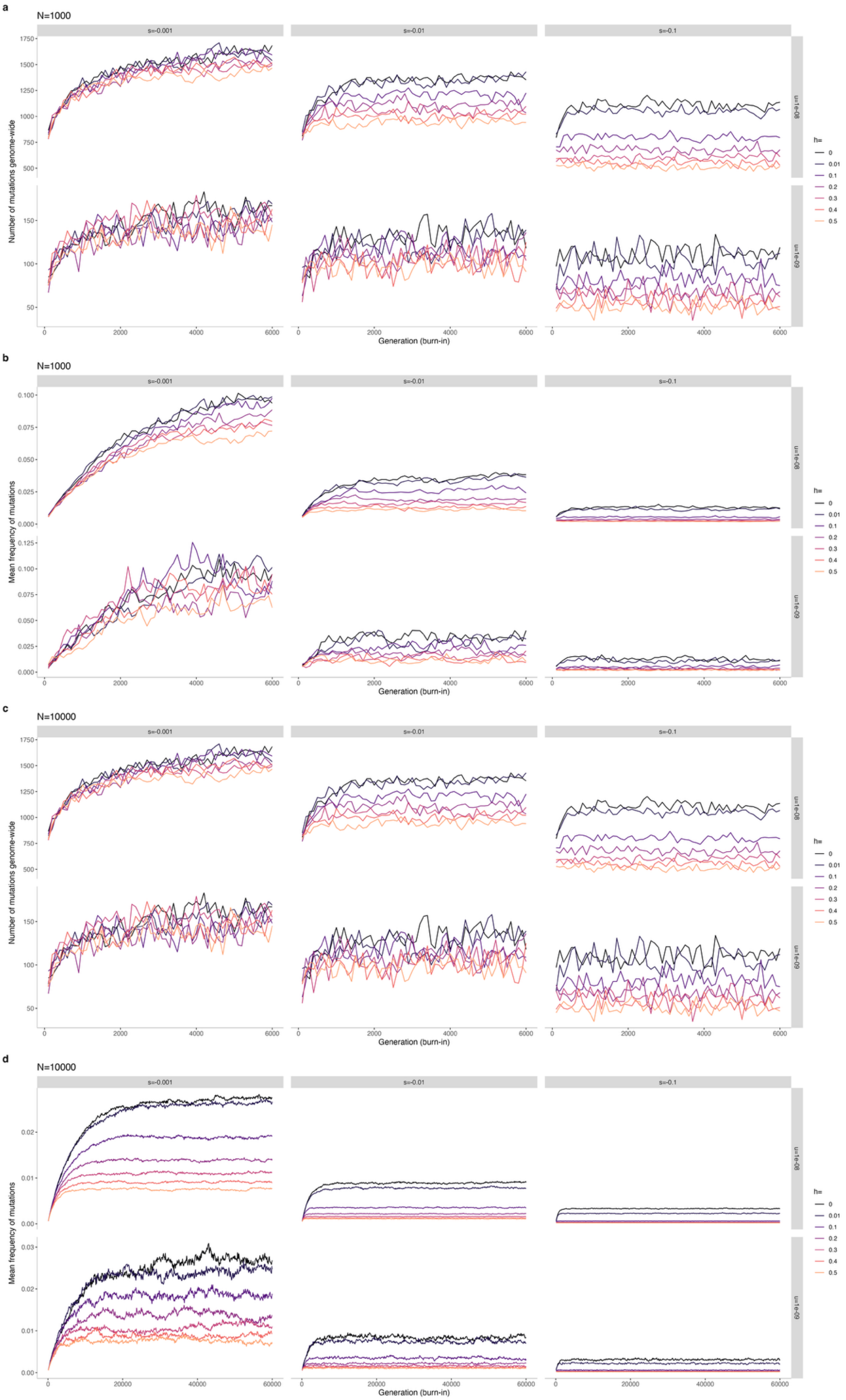
a
N=1000
s=-0.001
s=-0.01
s=-0.1
u=1e-08
u=1e-09
Number of mutations genome-wide
Generation (burn-in)
h=
0
0.01
0.1
0.2
0.3
0.4
0.5
b
N=1000
Mean frequency of mutations
c
N=10000
Number of mutations genome-wide
d
N=10000
Mean frequency of mutations

**Figure S14 | Population evolution during the burn-in phase.**

For the simulations of the effect of different parameters (notably *h* and *s*) on the fate of inversions, for each set of parameter values, one simulation was run and used as the initial state after a burn-in period of $6N$ generations (see methods). We display here the evolution of the population during the burn-in phase. Each trajectory represents the state of one simulation. **a,c** Number of mutations segregating in the genome. **b**,**d** Mean frequency of segregating mutations in the genome. This shows that each population had reached an equilibrium state at the end of the burn-in period.
